# Foliar application of mixed DAMP- and MAMP-oligosaccharides enhances root growth and fruit yield in okra and tomato in greenhouse experiments

**DOI:** 10.64898/2025.11.30.691324

**Authors:** Sreynich Pring, Akira Ashida, Yuki Shima, Yuki Iwamoto, Makoto Saito, Maurizio Camagna, Aiko Tanaka, Ikuo Sato, Sotaro Chiba, Daigo Takemoto

## Abstract

Plants perceive diverse endogenous and microbial molecular patterns that shape their defense, growth, and productivity. Damage-associated molecular patterns (DAMPs) such as cello- and xylo-oligosaccharides, and microbe-associated molecular patterns (MAMPs) such as chitin-oligosaccharides, are known to promote both development and immunity. Here, we evaluated the combined application of these three oligosaccharides (Oligo-Mix) in okra and cherry tomato under greenhouse conditions. Regular foliar treatment with Oligo-Mix enhanced root elongation and biomass accumulation, and increased fruit number and yield in both crops. RNA-seq analysis of Oligo-Mix-treated tomato leaves showed coordinated induction of photosynthetic genes, components of the translational machinery, cell-wall remodeling factors, and enzymes involved in flavonoid/anthocyanin biosynthesis, indicating broad enhancement of metabolic capacity and protective secondary pathways. Together, these results demonstrate that Oligo-Mix functions as a biostimulant that supports both vegetative growth and fruit production, offering a sustainable approach to improving crop performance.

## Introduction

Plants possess a sophisticated regulatory system that enables them to perceive and respond to a variety of external and internal molecules to ensure survival, growth, and productivity. Among these molecules, damage-associated molecular patterns (DAMPs) and microbe-associated molecular patterns (MAMPs) play key roles in regulating plant immunity, growth, and development (Boller and Felix 2009; Gust et al. 2017). DAMPs are endogenous molecules released upon cellular damage or stress, serving as alarm signals that activate defense mechanisms and tissue regeneration processes (De Lorenzo et al. 2019). Cello- and xylo-oligosaccharides (COS and XOS), derived from cellulose and hemicellulose in the plant cell wall, respectively, are well-known DAMPs (Souza et al. 2017; Claverie et al. 2018; Pring et al. 2023). In contrast, MAMPs are conserved microbial molecules that trigger plant defense mechanisms and facilitate beneficial interactions with rhizosphere microorganisms. Representative examples include bacterial flagellin (Gómez-Gómez and Boller 2000), peptidoglycan, a key structural component of bacterial cell walls (Gust et al. 2017), as well as chitin and β-glucan, which are major cell wall constituents of fungi (Sharp et al. 1984; Felix et al. 1993). In addition, 9-methyl-4,8-sphingadienine, a ceramide substructure present in fungi and oomycetes, as well as 5,8,11,14-tetraene-type fatty acids contained in microbial diacylglycerols, have been identified as MAMPs (Kato et al. 2022; Monjil et al. 2024). Some MAMPs are specific to certain groups of pathogenic microbes, such as elicitins secreted by oomycete pathogens (Derevnina et al. 2016) and fungal ethylene-inducing xylanases (EIXs) (Dean et al. 1991; Ron and Anvi 2004).

In sustainable agriculture, strategies to enhance plant growth and crop productivity while reducing chemical inputs have become a major focus. Recent studies have suggested that oligosaccharides derived from DAMPs and MAMPs can act as bioactive molecules, modulating root growth and development, enhancing nutrient uptake, and improving overall plant resilience (van Aubel et al. 2014). In another study, DAMPs cello- and xylo-oligosaccharide (COS and XOS) derived from plant cell wall and PAMPs chitosan oligosaccharide were shown to improve tomato seedling growth and chilling resistance (He et al. 2022). These oligosaccharides not only trigger immune responses but also stimulate growth-promoting pathways, leading to increased root biomass. However, although the individual effects of oligosaccharides have been investigated, their synergistic interactions in promoting plant growth and productivity remain largely understudied. In our previous research, a combination of DAMPs COS and XOS and MAMP chitin-oligosaccharides (CHOS), designated as Oligo-Mix, exerted positive effects on tomato growth and root development. Furthermore, in cucumber, Oligo-Mix enhanced seed germination and overall plant growth, along with increased chlorophyll content in the leaves (Pring et al. 2023). Moreover, it enhanced plant defenses against fungal pathogens such as powdery mildew and anthracnose disease (Pring et al. 2025a), suggesting that Oligo-Mix represents a promising biostimulant for agricultural applications. Biostimulants are characterized by two main functions: the first serves as a plant growth enhancer, and the second acts as a plant protection agent (Drobek et al. 2019; Jiang et al. 2024). Unlike fertilizers, biostimulants do not provide nutrients directly but instead enhance the plant’s capacity to utilize nutrients already present in the soil. Generally, their components are derived from natural substances of plants or microbial origin that improve crop vitality and condition without causing adverse effects (Calvo et al. 2014). By enhancing nutrient uptake, stimulating root development, and bolstering plant defenses, biostimulants are considered to play crucial roles in sustainable agriculture (Du Jardin 2015; Colla et al. 2017).

Tomato (*Solanum lycopersicum*) is one of the most important vegetable crops worldwide due to its high economic and nutritional value, whereas okra (*Abelmoschus esculentus*) is a regionally important crop cultivated extensively across tropical and subtropical regions (Elkhalifa et al. 2021). Okra is cultivated as a major crop in countries such as India and Nigeria, and it is also widely grown and commonly consumed in Japan, particularly in temperate to warm regions (Takahata et al., 2011). However, the productivity of these crops is often limited by environmental stress, soil nutrient deficiencies, and pathogen attacks (Akanbi et al. 2010; El-Saadony et al. 2022). Sustainable approaches that enhance root growth, nutrient absorption, and overall plant vigor are crucial for improving yield and ensuring food security. Oligosaccharides derived from DAMPs and MAMPs, as well as their effective combination, Oligo-Mix, offers a promising alternative to synthetic growth stimulants by activating natural plant signaling pathways and promoting beneficial rhizosphere interactions (Pring et al. 2023, 2025a).

Our previous studies demonstrated that Oligo-Mix promotes plant growth and disease resistance under controlled growth conditions (Pring et al., 2023, 2025a). In the present study, we extended these findings by applying Oligo-Mix to okra and tomato plants grown in a greenhouse, using weekly foliar spraying to assess its effects on growth promotion and productivity. Oligo-Mix effectively enhanced shoot and root development, including increases in root length and biomass, in both crops. Furthermore, Oligo-Mix treatment significantly increased fruit number and overall yield.

## Materials and Methods

### Plant materials and growth conditions

Seeds of okra (*Abelmoschus esculentus* cv. Green Sord, Takii Seed, Kyoto, Japan) and cherry tomato (*Solanum lycopersicum* cv. Pepe, Takii Seed) were sown in small pots containing a commercial soil mix (Sakata Super Mix A, Sakata Seed Corporation, Yokohama, Japan), and grown in a plant growth chamber under 16 h light / 8 h dark cycle at 19-28 °C, 76 % relative humidity. Two-week-old seedlings were transferred to larger pots (28 cm diameter × 22 cm height) filled with Sakata Super Mix A and grown in a greenhouse at the Togo field, Nagoya University (Togo, Aichi, Japan). Okra and tomato plants were irrigated twice daily using an automatic irrigation system installed in the greenhouse, without the addition of supplemental nutrients. Okra and tomato seeds were sown on 9 May 2024, and the seedlings transferred to larger pots were grown for additional 17 weeks (okra) and 23 weeks (tomato) for data collection.

### Experimental design

The experiment was conducted in the greenhouse and the layout was designed with two different treatments and three replicates, resulting in a total of six plots of okra and six plots of tomato. Each biological replicate consisted of 6 plants. The distance between plots was 110 cm x 50 cm, and plants were spaced 30 cm apart within each row. The arrangement of the experimental plots for tomato and okra is illustrated in Fig. 1. Each plot contained both Oligo-Mix treatment (O) and control (C) groups arranged alternately in three replicate rows, with six plants per row. The spacing between rows (1.1 m) and between plant lines (0.5 m) was maintained to minimize interference between treatments and to ensure uniform growth conditions.

**Fig. 1.**
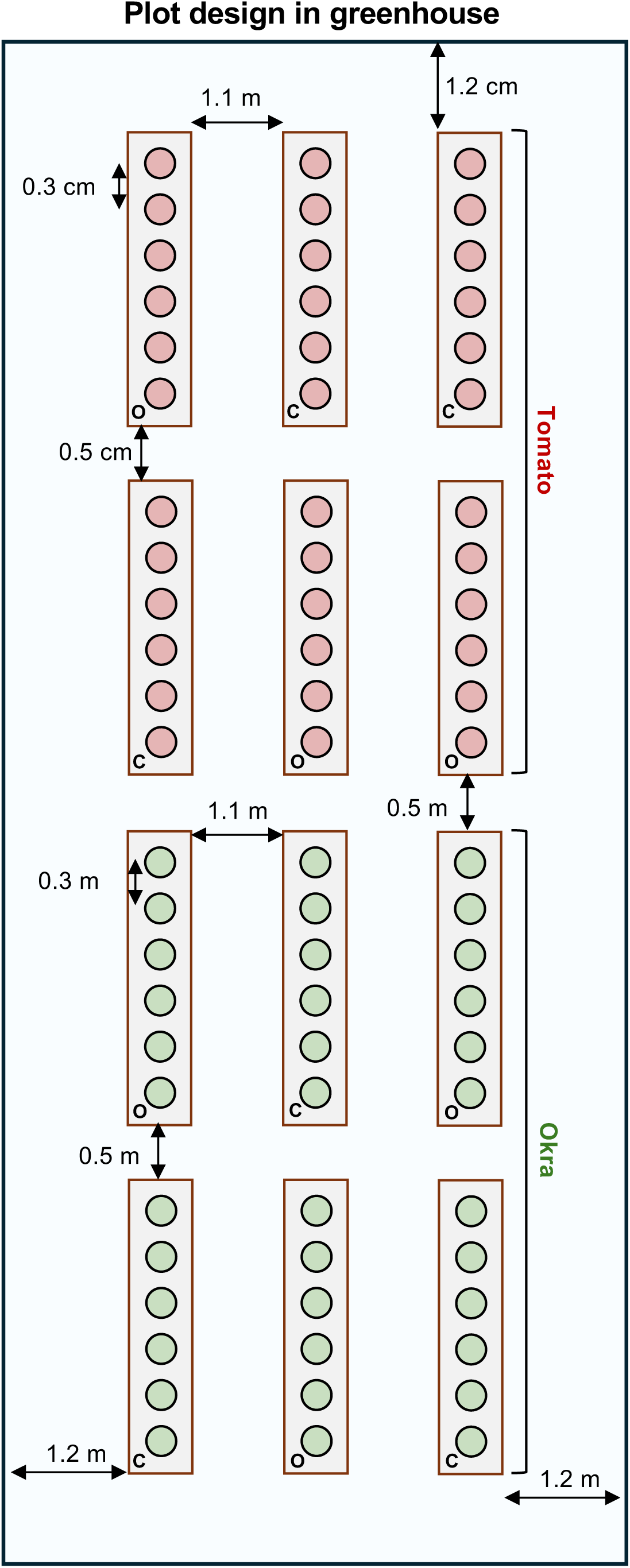
Experimental layout for greenhouse cultivation of tomato and okra plants. Tomato (*Solanum lycopersicum* cv. Pepe) and okra (*Abelmoschus esculentus* cv. Green Sord) plants were arranged in separate sections of the greenhouse. Each treatment consisted of three replicate rows containing six plants per row. Circles represent individual plants, with O indicating Oligo-Mix treatment and C indicating control plants. The spacing between rows and plants is shown in the diagram.

### Oligosaccharide mixture used in this study and the application

The purity and average degrees of polymerization (DP) of oligosaccharides used in this study were confirmed by gel permeation chromatography. The undiluted Oligo-Mix stock used in this study was composed of 20 mg/ml cello-oligosaccharide (COS, sourced from cotton linter, purity >99 %, average DP =3.4), 40 mg/ml xylo-oligosaccharide (XOS, from corn cob, purity >99 %, average DP = 2.7), and 20 mg/ml chitin-oligosaccharide (CHOS, from shrimp shell, purity >99 %, average DP =3.5) (Pring et al. 2023). Both Oligo-Mix and the control solutions contained water-soluble P (P_2_O_5_, 4.9% w/v) and K (K_2_O, 3.6% w/v). Oligo-Mix and control solution were diluted to 1:1000 and applied to plants by spraying. Oligo-Mix and control solutions were provided by Resonac Corporation (Tokyo, Japan). Five days after germination, each seedling was foliar-sprayed approximately 250 μl Oligo-Mix or control solution. Subsequently, 2-week-old seedlings were transplanted into pots and grown in a greenhouse, where they were continuously sprayed with the solution once per week (approximately 500 ml, for okra, and 730 ml for tomato per 18 plants) until sampling.

### Plant and root growth, and biomass measurement

Approximately 7- and 17-week-old okra plants and 23-week-old tomato plants were measured for shoot biomass, root length, and root biomass. Each plant was cut off above the ground, leaving the root portion intact. The fresh mass of the shoots was measured immediately after cutting. Roots were rinsed several times with water until all soil particles were completely removed. Root length was measured from the cutting point to the longest root. Subsequently, for dry mass determination, the shoot and root samples were placed in the greenhouse and air-dried for one week under summer temperatures ranging from 38 to 40 °C. Once fully dried, the dry biomass (g per plant) was measured.

### Yield quantity measurement

Yield was measured based on the quantification of fruit number and fruit weight per plant. Fruits of okra and tomato were harvested once a week after treatment. The harvested fruits were counted per plant, rinsed thoroughly with clean water, wiped with laboratory tissue paper, and then the fresh weight of each fruit was measured using a digital weighing scale. The average yield number per plant, and the average yield weight was calculated as follows:

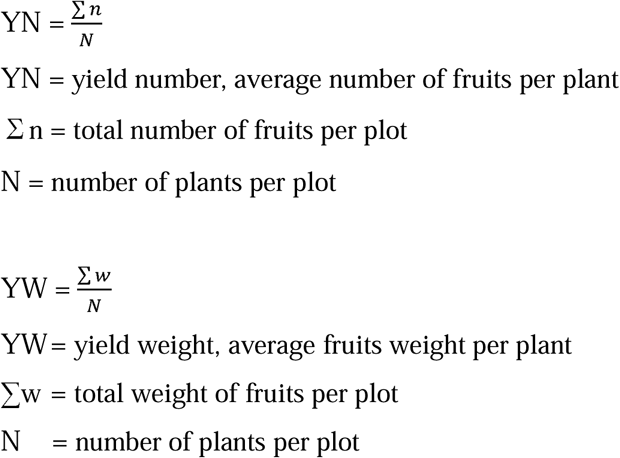

### RNA-seq and Statistical analysis

Tomato plants (cv. Renaissance, Sakata Seed, Japan) were grown in a growth-room under 16 h light / 8 h dark cycle at 23 °C. After germination of seedlings, plants were sprayed with water or Oligo-Mix from 3 days after the seedling germination and then once every week (4 times). Total RNA was extracted 24 h after last treatment using the RNeasy Plant Mini Kit (QIAGEN, Germany). Evaluation of RNA quality, library construction, and sequencing were performed as previously described (Kuroyanagi et al., 2022; Imano et al., 2023). Raw pair-end reads were quality-filtered and adapter-trimmed with fastp (v0.23.4, Chen et al., 2018; default settings with automatic adapter detection). Trimmed reads were aligned to the *Solanum lycopersicum* reference genome SLM_r2.1 (GCF_036512215.1) using HISAT2 (Kim et al., 2015) implemented in the HISAT2-pipeline (v1.0.8; single-end mode, default parameters, Camagna 2025), and transcript assembly and quantification were performed with StringTie (Pertea et al., 2015). Differential expression analysis was carried out using DESeq2 (v1.42.1, Love et al., 2014) with the design formula - condition. P values were adjusted for multiple testing using the Benjamini-Hochberg procedure, and genes with adjusted p-values (False discovery rate, FDR) < 0.05 and log_2_FC (fold change) > 1 were considered differentially expressed. For data visualization, MA plots were generated from DESeq2 results, and volcano plots were constructed using log_2_FC and −log_10_(P) values. Principal component analysis (PCA) was performed on rlog-transformed (or variance-stabilized) counts for all genes to assess global expression differences between treatments. RNA-seq data for tomato reported in this work are available in GenBank under the accession number DRA016704. Statistical analyses were performed using Student’s *t*-test, and SRplot (https://www.bioinformatics.com.cn/en), a free online data analysis and visualization platform, was used to generate the graphs.

## Results

### Oligo-Mix promotes root development and growth performance in okra

Oligo-Mix, a mixture of cello-oligosaccharide (COS), xylo-oligosaccharide (XOS), and chitin-oligosaccharide (CHOS), has been reported to promote plant and root growth in tomato (Pring et al. 2023) and cucumber (Pring et al. 2025a) under laboratory conditions. To further evaluate its effect on plant growth promotion and productivity under greenhouse conditions, two different plant species, okra (*Abelmoschus esculentus*, Malvaceae) and cherry tomato (*Solanum lycopersicum*, Solanaceae), were used in this study.

Seedlings of okra were sprayed with Oligo-Mix or a control solution 5 days after germination, transplanted into larger pots at 2 weeks, and then grown in the greenhouse, where they received weekly foliar applications. During the early growing stage, 7-week-old okra were harvested to measure plant height, root length, and both shoot and root biomass. The results demonstrated that Oligo-Mix treatment promoted a more fibrous root system and significantly increased root length compared to the control (Fig. 1A). Another eleven plants were retained until the final fruit harvest (17-week-old plants) for further evaluation, and similar improvements in plant and root growth were observed. As shown in Fig. 1B and C, Oligo-Mix treatment significantly enhanced root development, resulting in longer roots and higher root biomass compared with the control. Although aboveground (shoot) biomass did not differ significantly between treatments (Figs. 2C and S1A), okra plants treated with Oligo-Mix exhibited an overall trend of improved growth. These results suggest that Oligo-Mix enhances root growth, thereby improving the plants’ potential for nutrient and water uptake.

**Fig. 2.**
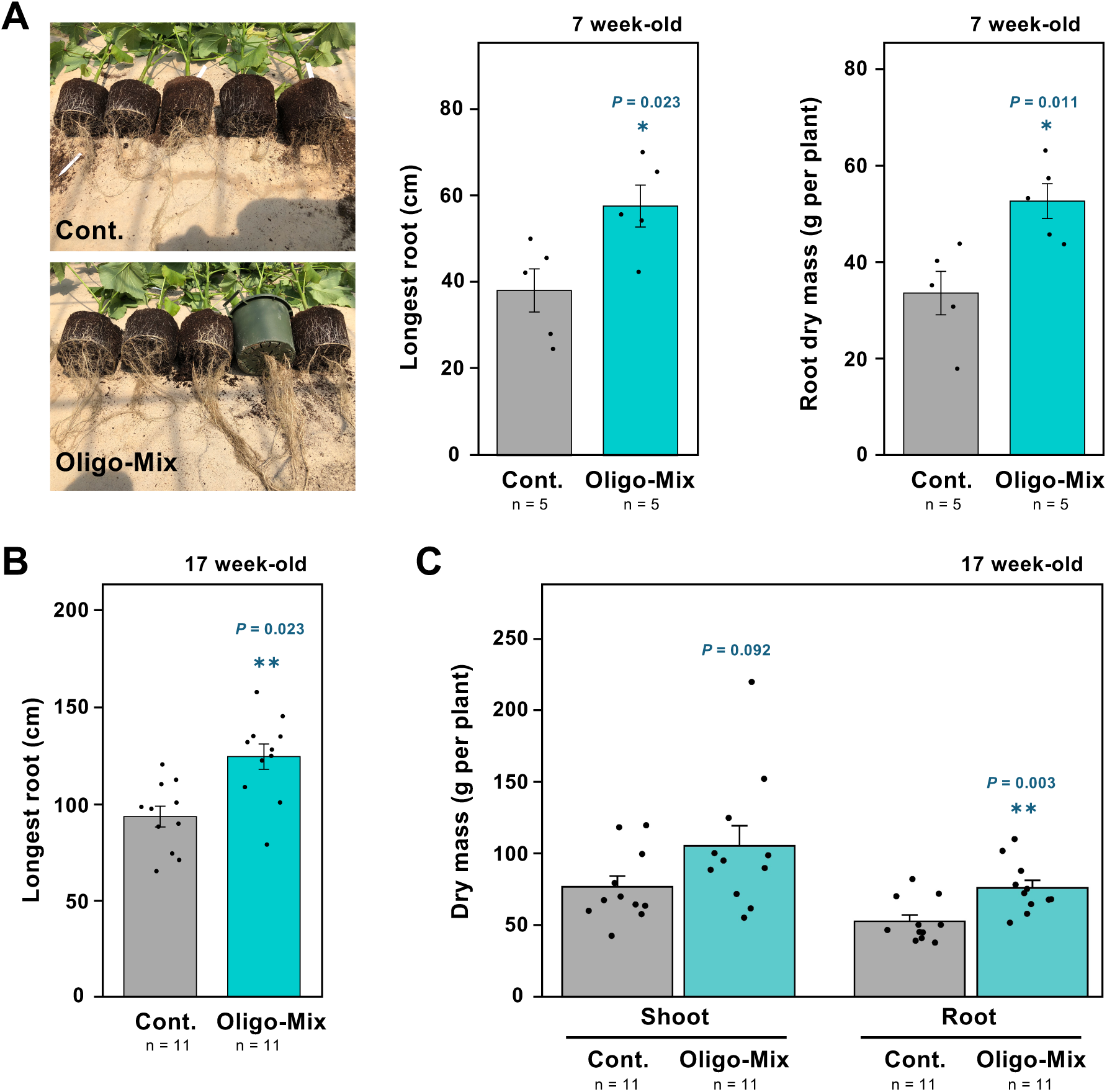
Effects of Oligo-Mix treatment on shoot and root growth in okra plants grown in the greenhouse. (**A)** Representative photographs show root growth of 7-week-old okra plants treated with the control solution (Cont.) or Oligo-Mix (20 μg/ml COS, 40 μg/ml XOS, and 20 μg/ml CHOS). The lengths of the longest roots and dry masses of roots per plant were measured. (n = 5). (**B)** The lengths of the longest roots of 17-week-old okra plants treated with control solution or Oligo-Mix were measured. (n = 11). **(C)** Dry masses of shoots (aboveground tissues) and roots per plant were measured for 17-week-old okra plants treated with the control solution or Oligo-Mix. (n = 11). Asterisks indicate a significant difference from control as assessed by a two-tailed Student’s *t*-test. * *p* < 0.05, ** *p* < 0.01.

### Oligo-Mix promotes fruit yield in okra plants

To determine whether enhanced root development translated into improved crop productivity, okra plants were grown under weekly foliar applications of Oligo-Mix or control solution until fruit maturation. Fruits were harvested once every week, totaling nine harvests throughout the growth period. The number of fruits per plant was recorded across all harvests (Fig. 3A). Oligo-Mix-treated plants consistently produced more fruits than the control at every harvest, resulting in a significantly higher average fruit number per plant across all nine harvests (Fig. 3B). A representative comparison of harvested fruits is shown in Fig. 3C. Analysis of fruit size distribution showed that Oligo-Mix increased fruit production in all weight categories: 5–15 g, 15–25 g, 25–35 g, and 35–45 g (Fig. 3D). Compared with the control, Oligo-Mix-treated plants produced 1.59-fold more small fruits (5–15 g), 1.12-fold more medium fruits (15–25 g), 1.14-fold more large fruits (25–35 g), and 1.71-fold more extra-large fruits (35–45 g). Together, these results indicate that Oligo-Mix enhances okra productivity primarily by increasing fruit number across all size categories, rather than by altering fruit size.

**Fig. 3.**
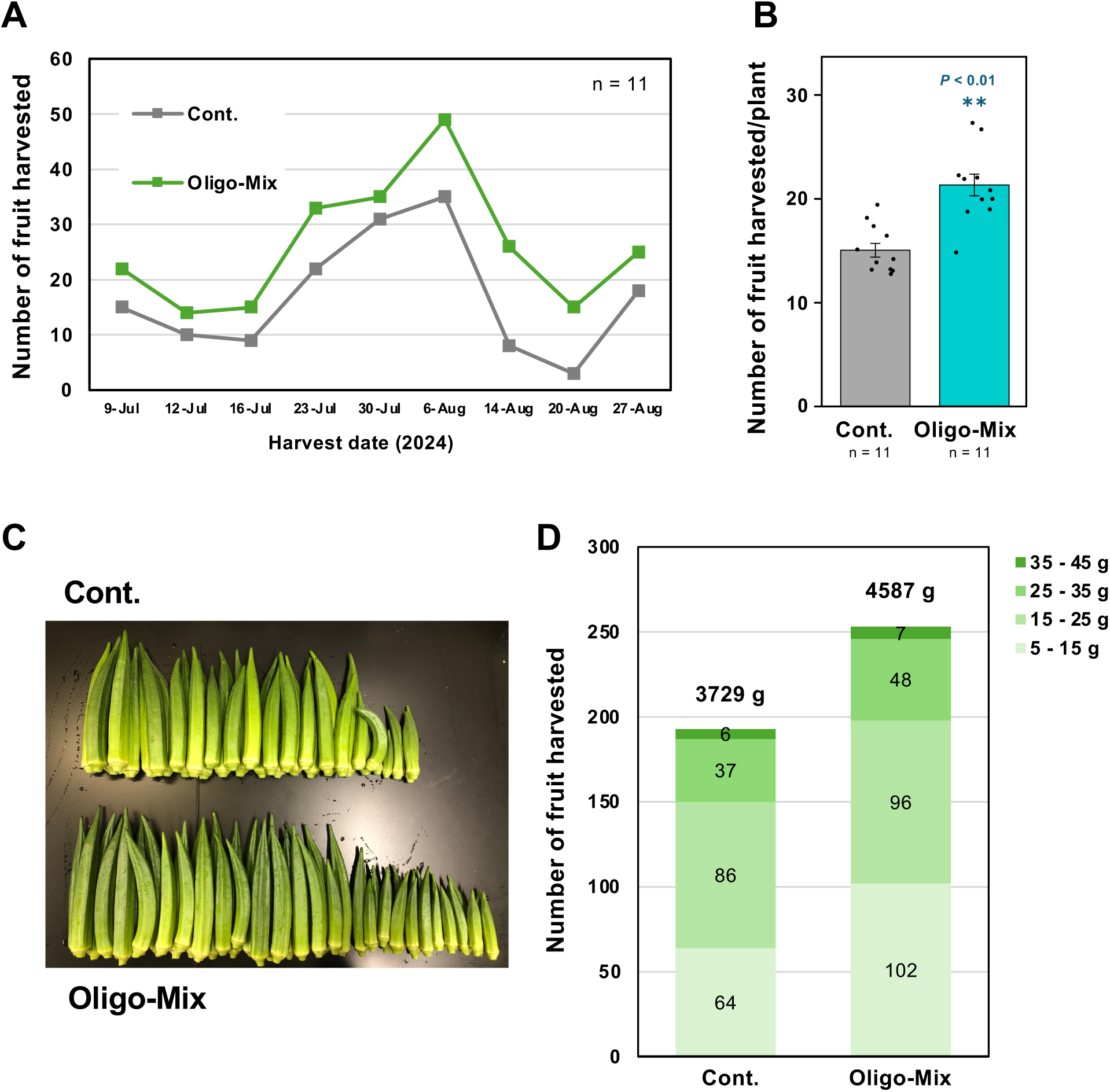
Effects of Oligo-Mix treatment on fruit yield of okra plants. **(A)** Changes in the number of harvested okra fruits across 11 plants over nine collection times for control (Cont.) and Oligo-Mix treatments. **(B)** Total number of okra fruits harvested per plant treated with the control solution or Oligo-Mix. Bars represent means ± SE (n = 11). Asterisks indicate a significant difference from control as assessed by a two-tailed Student’s *t*-test. ** p < 0.01. **(C)** Representative photograph of harvested okra fruits from control (top) and Oligo-Mix-treated (bottom) plants. **(D)** Distribution of all harvested okra fruits from 11 plants by weight class (5–15 g, 15–25 g, 25–35 g, and 35–45 g) for control and Oligo-Mix treatments. The total fruit weight per treatment is shown above each bar.

### Effect of Oligo-Mix on plant and root growth of cherry tomato

For cherry tomato plants, approximately 23-week-old individuals were evaluated for plant and root growth at the end of the cultivation period. Representative root systems are shown in Fig. 4A. Because of the extended growth period, parts of the root system often remained embedded in the soil and could not be fully recovered. As a result, Oligo-Mix–treated plants showed a trend toward longer roots (Fig. 4B), although the difference was not statistically significant. In contrast, root dry biomass was significantly higher in Oligo-Mix–treated plants compared with the control (Fig. 4C), indicating enhanced belowground biomass accumulation. Together with the results obtained in okra, these findings demonstrate that Oligo-Mix consistently promotes root biomass production across different crop species and growth durations.

**Fig. 4.**
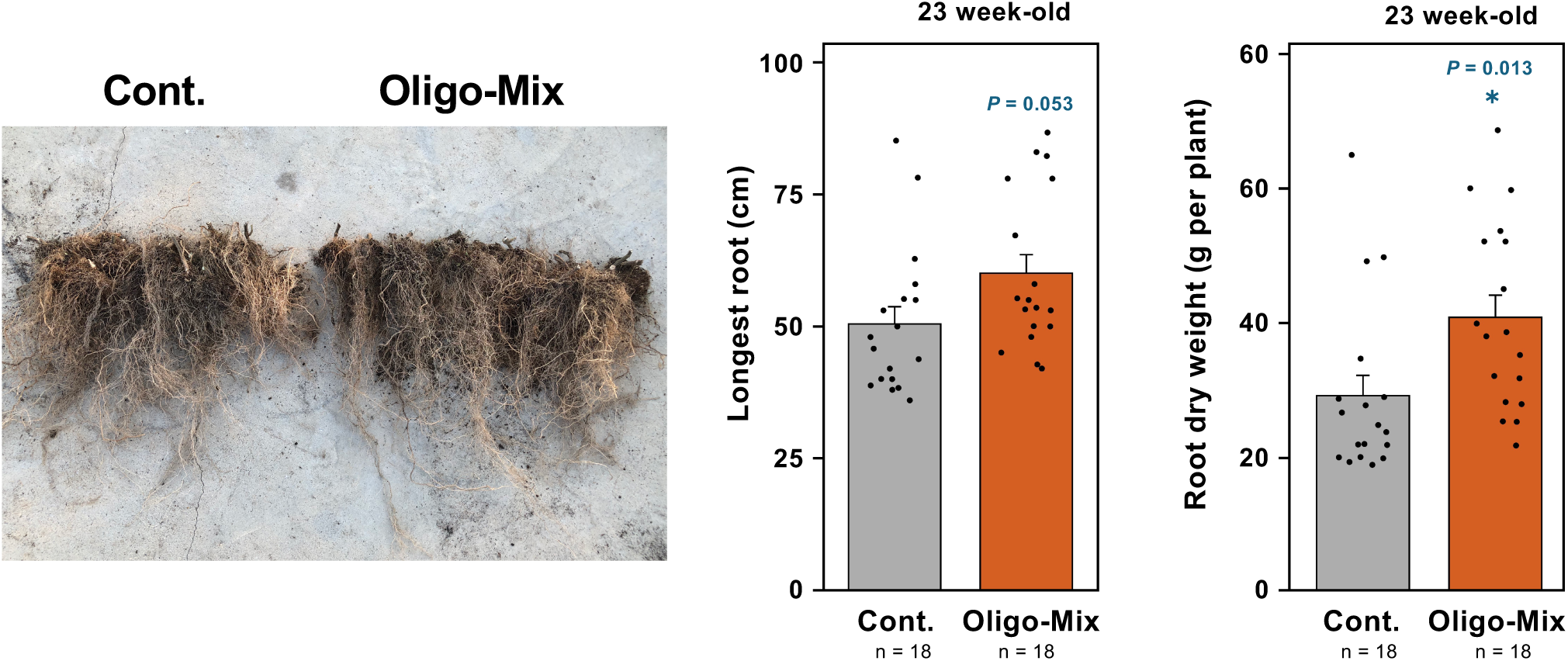
Effects of Oligo-Mix treatment on root growth in cherry tomato plants grown in the greenhouse. (Left) Representative photographs show root growth of 23-week-old cherry tomato plants treated with the control solution (Cont.) or Oligo-Mix (20 μg/ml COS, 40 μg/ml XOS, and 20 μg/ml CHOS). (Middle) The lengths of the longest roots and (Right) dry masses of roots per plant were measured. Asterisks indicate significant difference from control as assessed by a two-tailed Student’s *t*-test. (n = 18) * *p* < 0.05.

### Oligo-Mix enhances fruit yield in cherry tomato plants

To further investigate the effect of Oligo-Mix on fruit yield, cherry tomato plants were grown under the same greenhouse conditions and treated weekly with either Oligo-Mix or control solution until fruit maturity. Fruits were harvested five times during the growing season (Fig. 5A). The average number of fruits per plant was calculated (Fig. 5B). While individual harvests showed no significant differences between treatments, the total average fruit number per plant across all harvests was significantly higher in Oligo-Mix–treated plants compared with the control (Fig. 5B). A representative comparison of harvested fruits, showing the greater fruit number in Oligo-Mix–treated plants is presented in Fig. 5C. The total fruit weight per plant also showed a significant increase following Oligo-Mix treatment (Fig. 5B). Analysis of fruit size distribution revealed that Oligo-Mix increased fruit production across all weight categories defined in Fig. 5D: < 5 g, 5-10 g, 10–15 g, and > 15 g. Compared with the control, Oligo-Mix-treated plants produced 1.36-fold more fruits < 5 g, 1.12-fold more fruits 5-10 g, 1.16-fold more fruits 10-15 g, and 1.54-fold more fruits > 15 g.

**Fig. 5.**
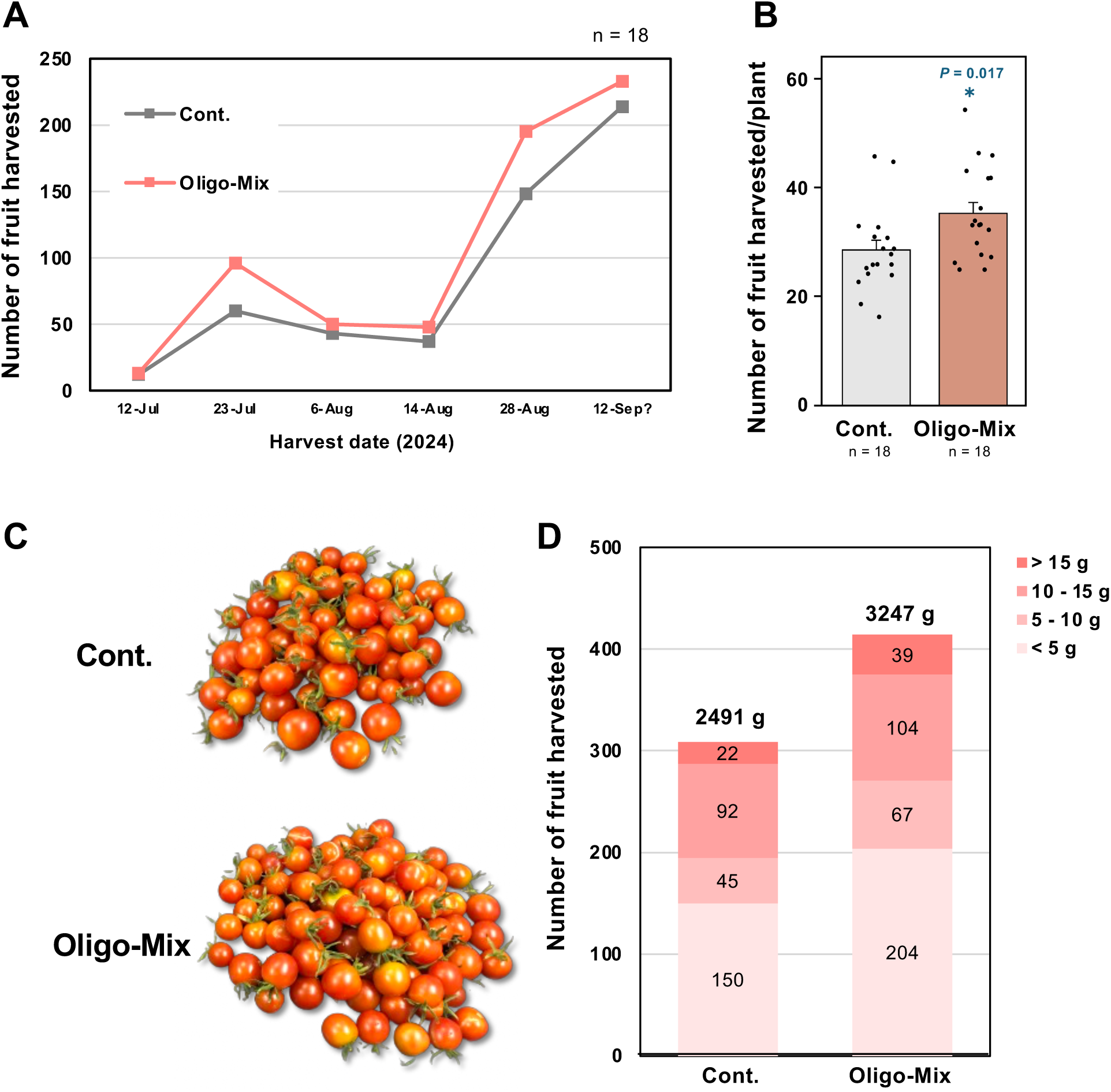
Effects of Oligo-Mix treatment on fruit yield of cherry tomato plants. **(A)** Changes in the number of harvested cherry tomato fruits across 18 plants over nine collection times for control (Cont.) and Oligo-Mix treatments. **(B)** Total number of cherry tomato fruits harvested per plant treated with the control solution or Oligo-Mix. Bars represent means ± SE (n = 18). Asterisks indicate a significant difference from control as assessed by a two-tailed Student’s *t*-test. * p < 0.05. **(C)** Representative photograph of harvested cherry tomato fruits from control (top) and Oligo-Mix-treated (bottom) plants. **(D)** Distribution of all harvested cherry tomato fruits from 18 plants by weight class (< 5 g, 5–10 g, 10–15 g, and >15 g) for control and Oligo-Mix treatments. The total fruit weight per treatment is shown above each bar.

Taken together with the results in okra, these findings indicate that Oligo-Mix enhances the capacity of plants to produce a greater number of fruits in multiple crop species. The synergistic combination of plant-derived oligosaccharides (COS and XOS) and a pathogen-derived oligosaccharide (CHOS) likely contributes to improved crop yield.

### Oligo-Mix enhances photosynthetic and stress-associated transcriptional programs in tomato

An expanded analysis of a previously obtained RNA-seq dataset from Oligo-Mix–treated tomato leaves (Pring et al., 2023), re-mapped to the updated tomato genome assembly SLM_r2.1 (Nagasaki et al., 2024), was performed to clarify the transcriptional changes underlying the physiological responses observed in this study. Oligo-Mix treatment induced a broad transcriptional reprogramming in tomato leaves (Table S1). The set of upregulated genes spanned multiple functional categories, including photosynthetic energy conversion, ribosome and RNA metabolism, cell-wall remodeling, and flavonoid/anthocyanin biosynthesis.

A prominent signature of the Oligo-Mix response was the consistent activation of photosynthesis-related genes. These included genes for photosystem II subunits (e.g., *psbT*, *psbI* and *psbZ*), the Ribulose-1,5-bisphosphate carboxylase/oxygenase large subunit (*rbcL*), Photosystem I assembly factor *ycf3*, ATP synthase CF0 subunit I (*atpF*), and the cytochrome b6-f complex subunit (Table S1 and Fig. 5A). In addition, a large number of genes including chloroplast ribosomal proteins (e.g., *rps14*, *rps19* and *rps3*), chloroplast tRNAs (e.g., tRNA-Ser and tRNA-Arg), the chloroplast *RNA polymerase* α*-subunit*, and *RNA pseudouridine synthase 6* were highly induced (Fig. 5B). These transcriptional and post-transcriptional components collectively indicate enhanced chloroplast translation capacity and RNA maturation. Their coordinated induction suggests that Oligo-Mix strengthens photosynthetic function not only through photosystem assembly and electron transfer, but also through accelerated biogenesis and functional maintenance of the chloroplast translational machinery. Genes involved in cell-wall dynamics were also represented among the Oligo-Mix-induced transcripts. These included genes for *glycine-rich cell wall protein*, two xyloglucan endotransglucosylase/hydrolase genes (*XTH32* and *XTH6*), a *pectinesterase/pectinesterase inhibitor*, *GPI-anchored lipid transfer protein* and *Lignin-forming anionic peroxidase* (Table S1 and Fig. 5C). Together, these proteins act on major components of the apoplast-including xyloglucan, pectin, and hydroxyproline-rich glycoproteins indicating active remodeling of the extracellular matrix. Such structural changes are likely to increase tissue flexibility, alter apoplastic transport properties, and enhance the plant’s readiness to respond to biotic or abiotic stress.

Genes involved in flavonoid and anthocyanin biosynthesis were also induced by Oligo-Mix. Core pathway enzymes, chalcone synthase (*CHS2*), chalcone–flavonone isomerase (*CHI1*), flavanone 3-dioxygenase (*F3H*), dihydroflavonol 4-reductase (*DFR*), and anthocyanidin synthase (*ANS*), were all upregulated, together with genes for downstream modification enzymes such as UDP-glycosyltransferases (e.g. *UGT75C1* and *UGT86A1*), anthocyanidin 3-O-glucosyltransferase, and anthocyanin acyltransferase (Table S1 and Fig. 5D). This coordinated induction of both core and tailoring steps indicates that Oligo-Mix enhances flavonoid metabolic flux, likely contributing to increased antioxidant capacity and stress tolerance.

In addition to these pathway-specific responses, Oligo-Mix broadly activated regulatory and signaling pathways, including factors involved in transcriptional control, hormone responses, and cellular stress signaling. The concurrent induction of genes related to chloroplast function, protein synthesis, secondary metabolism, and defense processes indicates a coordinated, system-wide adjustment of cellular physiology. Together, these transcriptional changes suggest that Oligo-Mix promotes a primed metabolic and defensive state that enhances the plant’s overall stress tolerance and growth potential.

## Discussion

This study demonstrated that foliar application of Oligo-Mix, a mixture composed of DAMPs (cello-oligosaccharides, COS; and xylo-oligosaccharides, XOS) together with a MAMP (chitin-oligosaccharides, CHOS), significantly enhanced root growth and increased fruit yield in both okra and cherry tomato under greenhouse conditions. These results are consistent with, and extend, the findings of Pring et al. (2023, 2025a), which showed that Oligo-Mix promoted shoot and root growth of tomato and cucumber under controlled laboratory environments. In addition, Oligo-Mix treatment conferred resistance in cucumber against fungal pathogens, including powdery mildew (*Podosphaera xanthii*) and anthracnose (*Colletotrichum orbiculare*) (Pring et al. 2025a). Taken together, these findings indicate that Oligo-Mix acts as a promising biostimulant that promotes plant growth and disease resistance across different species.

In Arabidopsis, the perception mechanisms for plant- and microbe-derived oligosaccharides have been well characterized. CHOS are recognized by the CERK1–LYK5 receptor complex (Miya et al. 2007; Cao et al. 2014), whereas COS are sensed through CORK1 (AT1G56145) and its related leucine-rich repeat-malectin receptor kinases AT1G56130 and AT1G56140, which function independently of the CERK1 pathway (Tseng et al. 2022; Martín-Dacal et al., 2023). Despite these distinct receptor systems, both types of oligosaccharides induce similar defense responses, including rapid reactive oxygen species (ROS) generation, MAP kinase phosphorylation, and activation of the *WRKY33* promoter, suggesting a convergence of signaling networks that link cell-damage perception with immune activation (Kawasaki et al. 2017; Claverie et al. 2018; Pring et al. 2023). DAMPs such as COS and XOS act as endogenous elicitors of tissue repair and regeneration, whereas CHOS, a fungal MAMP, triggers immune signaling. RNA-seq of Arabidopsis seedlings revealed broad transcriptional activation by each oligosaccharide, with the combined Oligo-Mix producing additive effects on gene expression, consistent with the priming of both developmental and defense networks (Pring et al., 2023).

The transcriptional changes observed in tomato leaves indicate that Oligo-Mix elicits a coordinated adjustment of multiple metabolic processes rather than acting on a single pathway. The simultaneous induction of photosynthesis-related genes, translational machinery, cell-wall remodeling factors, and flavonoid/anthocyanin enzymes points to an integrated shift linking primary and secondary metabolism. Enhanced carbon assimilation, increased protein synthesis capacity, and strengthened structural and antioxidant systems appear to act together to support growth and stress adaptation. Within this context, activation of flavonoid and anthocyanin biosynthesis likely contributes antioxidative and photoprotective functions, helping buffer ROS generated under elevated photosynthetic activity and stabilizing chloroplast performance. These observations suggest that Oligo-Mix modulates a broader regulatory landscape that couples resource acquisition with protective metabolism, providing a mechanistic basis for the enhanced root development and yield observed in this study.

Consistent with this model, RNA-seq analysis of tomato leaves treated with Oligo-Mix showed a clear trend toward increased expression of photosynthesis-related genes (Fig. 6; Pring et al., 2023). Similar enhancement of chlorophyll content and photosynthetic performance was reported in cucumber (Pring et al., 2025a), supporting the idea that Oligo-Mix stimulates carbon assimilation. In okra, Oligo-Mix-treated plants exhibited significantly longer roots and greater dry biomass, while shoot biomass remained comparable to controls (Fig. 2), suggesting a preferential allocation of assimilates to belowground tissues. However, it remains unclear whether these growth-promoting effects arise primarily through foliar signaling or through secondary effects mediated in the rhizosphere; further study will be required to disentangle these mechanisms.

**Fig. 6.**
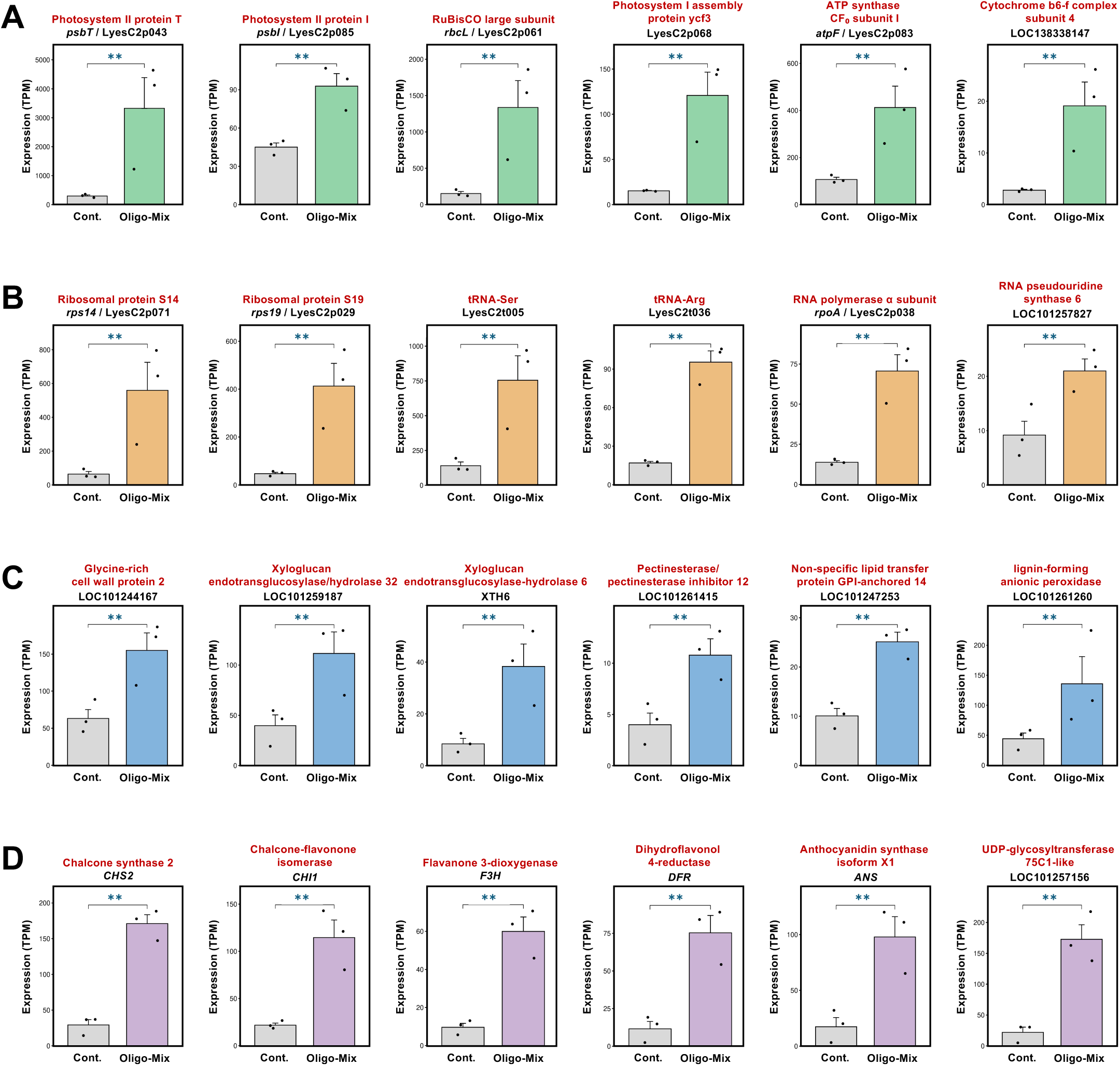
Expression profiles of representative tomato genes upregulated by Oligo-Mix treatment. Transcript levels (TPM values) were obtained from RNA-seq analysis of tomato leaves treated with control solution (Cont.) or Oligo-Mix (20 µg/mL COS, 40 µg/mL XOS, 20 µg/mL CHOS) for 24 h. **(A)** Genes involved in photosynthesis, including components of Photosystem II, Rubisco, and ATP synthase. **(B)** Genes involved in protein synthesis such as chloroplast ribosomal proteins, tRNAs, and RNA-processing factors. **(C)** Genes related to modification and strengthening of the cell wall, including xyloglucan-remodeling enzymes and peroxidases. **(D)** Genes involved in producing and modifying flavonoids and anthocyanins, including key biosynthetic enzymes and related transferases. Asterisks indicate significant differences compared with the control (\*\**FDR* < 0.01). See Table S1 for the complete list of Oligo-Mix-induced genes.

In addition, Oligo-Mix induced genes involved in flavonoid and anthocyanin biosynthesis (Fig. 6 and Table S1). These metabolites function as key antioxidants and photoprotective compounds that mitigate ROS accumulation, stabilize photosynthetic machinery, and contribute to stress tolerance (Agati et al. 2012; Nakabayashi and Saito 2015). Flavonoids also modulate auxin transport and can influence root architecture (Brown et al. 2001). Thus, the enhanced activation of flavonoid pathways observed in this study likely provides both antioxidative buffering and developmental modulation, supporting the improved growth and stress-related traits associated with Oligo-Mix treatment.

In summary, this study demonstrates that Oligo-Mix enhances root development and increases fruit yield in both okra and tomato under greenhouse conditions, thereby validating and extending previous reports obtained in other species under controlled environments. The yield gains observed here, achieved without additional fertilizer input, highlight the potential of Oligo-Mix as a sustainable biostimulant. These coordinated physiological effects suggest that Oligo-Mix supports a balanced activation of growth- and defense-related processes. Future studies should investigate how oligosaccharide perception interacts with hormonal signaling and resource allocation, and assess the robustness of these effects under field conditions across diverse crop species.

## Acknowledgements

This work was supported partially by a Grant-in-Aid for Scientific Research (B) (23H02212) and a Grant-in-Aid for JSPS Research Fellow (23KF0146) from the Japan Society for the Promotion of Science (JSPS), Aichi Agricultural Innovation Project 2026 (Aichi prefecture, Japan) and donation from Resonac Corporation (former Showa Denko K.K., Tokyo, Japan).

## Author Contributions

DT designed the research. SP and YI conducted the experiments. AA, YS, AT, and DT analyzed the data. Funding was acquired by DT. MS provided materials. AA, CM, IS, SC, and DT supervised the experiments. SP and DT wrote the original manuscript, and DT edited the manuscript.

## Data Availability

The data will be made available upon request from the authors.

## References

Agati G, Azzarello E, Pollastri S, Tattini M (2012) Flavonoids as antioxidants in plants: Location and functional significance. Plant Sci 196: 67–76. 10.1016/j.plantsci.2012.07.014

Akanbi WB, Togun, AO, Adediran JA, Adediran JA, Ilupeju EAO (2010) Growth, dry matter and fruit yields components of okra under organic and inorganic sources of nutrient. Am-Euras J Sustain Agric 4:1–13.

Boller T, Felix GA (2009) Renaissance of elicitors: perception of microbe-associated molecular patterns and danger signals by pattern-recognition receptors. Annu Rev Plant Biol. 60:379–406. 10.1146/annurev.arplant.57.032905.105346

Brown DE, Rashotte AM, Murphy AS, Normanly J, Tague BW, Peer WA, Taiz L, Muday GK (2001) Flavonoids act as negative regulators of auxin transport in vivo in Arabidopsis. Plant Physiol 126: 524–535. 10.1104/pp.126.2.524

Claverie J, Balacey S, Lemaître-Guillier C, Brulé D, Chiltz A, Granet L, Noirot E, Daire X, Darblade B, Héloir MC, Poinssot B (2018) The cell wall-derived xyloglucan is a new DAMP triggering plant immunity in *Vitis vinifera* and *Arabidopsis thaliana*. Front Plant Sci 9:1725. 10.3389/fpls.2018.01725

Calvo P, Nelson L, and Kloepper JW (2014) Agricultural uses of plant biostimulants. Plant Soil 383:3–41. 10.1007/s11104-014-2131-8

Camagna M (2025) HISAT2-pipeline. Zenodo. https://zenodo.org/records/16902402

Chen S, Zhou Y, Chen Y, Gu J (2018) fastp: an ultra-fast all-in-one FASTQ preprocessor. Bioinformatics 34: i884–i890. 10.1093/bioinformatics/bty560

Colla G, Hoagland L, Ruzzi M, Cardarelli M, Bonini P, Canaguier R, Rouphael Y (2017) Biostimulant action of protein hydrolysates: unraveling their effects on plant physiology and microbiome. Front Plant Sci 8:2202. 10.3389/fpls.2017.02202

De Lorenzo G, Ferrari S, Giovannoni M, Mattei B, Cervone F. (2019) Cell wall traits that influence plant development, immunity, and bioconversion. Plant J 97:134–147. 10.1111/tpj.14196

Dean JF, Gross KC, Anderson JD. (1991) Ethylene biosynthesis-inducing xylanase : III. Product characterization. Plant Physiol 96:571–6. 10.1104/pp.96.2.571

Derevnina L, Dagdas YF, De la Concepcion JC, Bialas A, Kellner R, Petre B, Domazakis E, Du J, Wu CH, Lin X, Aguilera-Galvez C, Cruz-Mireles N, Vleeshouwers VG, Kamoun S (2016) Nine things to know about elicitins. New Phytol 212:888–895. 10.1111/nph.14137

Drobek M, Frąc M, Cybulska J (2019) Plant biostimulants: Importance of the quality and yield of horticultural crops and the improvement of plant tolerance to abiotic stress - A review. Agronomy 9:335. 10.3390/agronomy9060335

Du Jardin P (2015) Plant biostimulants: Definition, concept, main categories and regulation. Sci Hortic 196:3–14. 10.1016/j.scienta.2015.09.021

El-Saadony MT, Saad AM, Soliman SM, Salem HM, Ahmed AI, Mahmood M, El-Tahan AM, Ebrahim AAM, Abd El-Mageed TA, Negm SH, Selim S, Babalghith AO, Elrys AS, El-Tarabily KA, AbuQamar SF (2022) Plant growth-promoting microorganisms as biocontrol agents of plant diseases: Mechanisms, challenges and future perspectives. Front Plant Sci 13:923880. 10.3389/fpls.2022.923880

Felix G, Regenass M, Boller T (1993) Specific perception of subnanomolar concentrations of chitin fragments by tomato cells: induction of extracellular alkalinization, changes in protein phosphorylation, and establishment of a refractory state. Plant J 4:307–316. 10.1046/j.1365-313X.1993.04020307.x

Gómez-Gómez L, Boller T (2000) FLS2: an LRR receptor-like kinase involved in the perception of the bacterial elicitor flagellin in Arabidopsis. Mol Cell 5:1003–1011. 10.1016/s1097-2765(00)80265-8

Gust AA, Pruitt R, Nürnberger T (2017) Sensing danger: Key to activating plant immunity. Trends Plant Sci 22:779–791. 10.1016/j.tplants.2017.07.005

He J, Han W, Wang J, Qian Y, Saito M, Bai W, Song J, and Lv G (2022) Functions of oligosaccharides in improving tomato seeding growth and chilling resistance. J Plant Growth Regul 41:535–545. 10.1007/s00344-021-10319-0

Imano S, Fushimi M, Camagna M, Tsuyama-Koike A, Mori H, Ashida A, Tanaka A, Sato I, Chiba S, Kawakita K, Ojika M, Takemoto D (2022) AP2/ERF transcription factor NbERF-IX-33 is involved in the regulation of phytoalexin production for the resistance of *Nicotiana benthamiana* to *Phytophthora infestans*. Front Plant Sci. 12:821574. 10.3389/fpls.2021.821574

Jiang Y, Yue Y, Wang Z, Lu C, Yin Z, Li Y, Ding X (2024) Plant biostimulant as an environmentally friendly alternative to modern agriculture. J Agric Food Chem 72:5107–5121. 10.1021/acs.jafc.3c09074

Kato H, Nemoto K, Shimizu M, Abe A, Asai S, Ishihama N, Matsuoka S, Daimon T, Ojika M, Kawakita K, Onai K, Shirasu K, Yoshida M, Ishiura M, Takemoto D, Takano Y, Terauchi R (2022) Recognition of pathogen-derived sphingolipids in Arabidopsis. Science 376:857–860. 10.1126/science.abn0650

Kim D, Langmead B, Salzberg SL (2015) HISAT: a fast spliced aligner with low memory requirements. Nat Methods 12: 357–360. 10.1038/nmeth.3317

Kuroyanagi T, Bulasag AS, Fukushima K, Ashida A, Suzuki T, Tanaka A, Camagna M, Sato I, Chiba S, Ojika M, Takemoto D (2022) *Botrytis cinerea* identifies host plants via the recognition of antifungal capsidiol to induce expression of a specific detoxification gene. PNAS Nexus1:pgac274. 10.1093/pnasnexus/pgac274

Love MI, Huber W, Anders S (2014) Moderated estimation of fold change and dispersion for RNA-seq data with DESeq2. Genome Biol 15: 550. 10.1186/s13059-014-0550-8

Monjil MS, Kato H, Ota S, Matsuda K, Suzuki N, Tenhiro S, Tatsumi A, Pring S, Miura A, Camagna M, Suzuki T, Tanaka A, Terauchi R, Sato I, Chiba S, Kawakita K, Ojika M, Takemoto D (2024) Two structurally different oomycete lipophilic microbe-associated molecular patterns induce distinctive plant immune responses. Plant Physiol 196:479–494. 10.1093/plphys/kiae255

Nagasaki H, Shirasawa K, Hoshikawa K, Isobe S, Ezura H, Aoki K, Hirakawa H (2024) Genomic variation across distribution of Micro-Tom, a model cultivar of tomato (*Solanum lycopersicum*). DNA Res 31:dsae016. 10.1093/dnares/dsae016

Nakabayashi R, Saito K (2015) Integrated metabolomics and transcriptomics for understanding stress responses in plants. Curr Opin Plant Biol 24: 10–16. 10.1016/j.pbi.2015.01.003

Pertea M, Pertea GM, Antonescu CM, Chang TC, Mendell JT, Salzberg SL (2015) StringTie enables improved reconstruction of a transcriptome from RNA-seq reads. Nat Biotechnol 33: 290–295. 10.1038/nbt.3122

Pring S, Kato H, Imano S, Camagna M, Tanaka A, Kimoto H, Chen P, Shrotri A, Kobayashi H, Fukuoka A, Saito M, Suzuki T, Terauchi R, Sato I, Chiba S, Takemoto D (2023) Induction of plant disease resistance by mixed oligosaccharide elicitors prepared from plant cell wall and crustacean shells. Physiol Plant 175:e14052. 10.1111/ppl.14052

Pring S, Kato H, Taniuchi K, Camagna M, Saito M, Tanaka A, Merritt BA, Argüello-Miranda O, Sato I, Chiba S, Takemoto D (2025a) Mixed DAMP/MAMP oligosaccharides promote both growth and defense against fungal pathogens of cucumber. Plant Sci 359:112578. 10.1016/j.plantsci.2025.112578

Ron M, Avni A (2004) The receptor for the fungal elicitor ethylene-inducing xylanase is a member of a resistance-like gene family in tomato. Plant Cell 16:1604–1615. 10.1105/tpc.022475

Sharp JK, Valent B, Albersheim P (1984) Purification and partial characterization of a β-glucan fragment that elicits phytoalexin accumulation in soybean. J Biol Chem 259:11312–11320.

Souza CA, Li S, Lin AZ, Boutrot F, Grossmann G, Zipfel C, Somerville SC (2017) Cellulose-derived oligomers act as damage-associated molecular patterns and trigger defense-like responses. Plant Physiol 173:2383–2398. 10.1104/pp.16.01680

Takahata K, Mine Y, Miura H (2011) Optimum emergence temperature for okra seeds. Trop Agric Dev 55:93–96. 10.11248/jsta.55.93

van Aubel G, Buonatesta R, Van Cutsem P (2014). COS-OGA: a novel oligosaccharidic elicitor that protects grapes and cucumbers against powdery mildew. Crop Prot 65:129–137. 10.1016/j.cropro.2014.07.015

