## Supplementary material for "Foliar application of mixed DAMP- and MAMP-oligosaccharides enhances root growth and fruit yield in okra and tomato in greenhouse experiments": Fig. S1

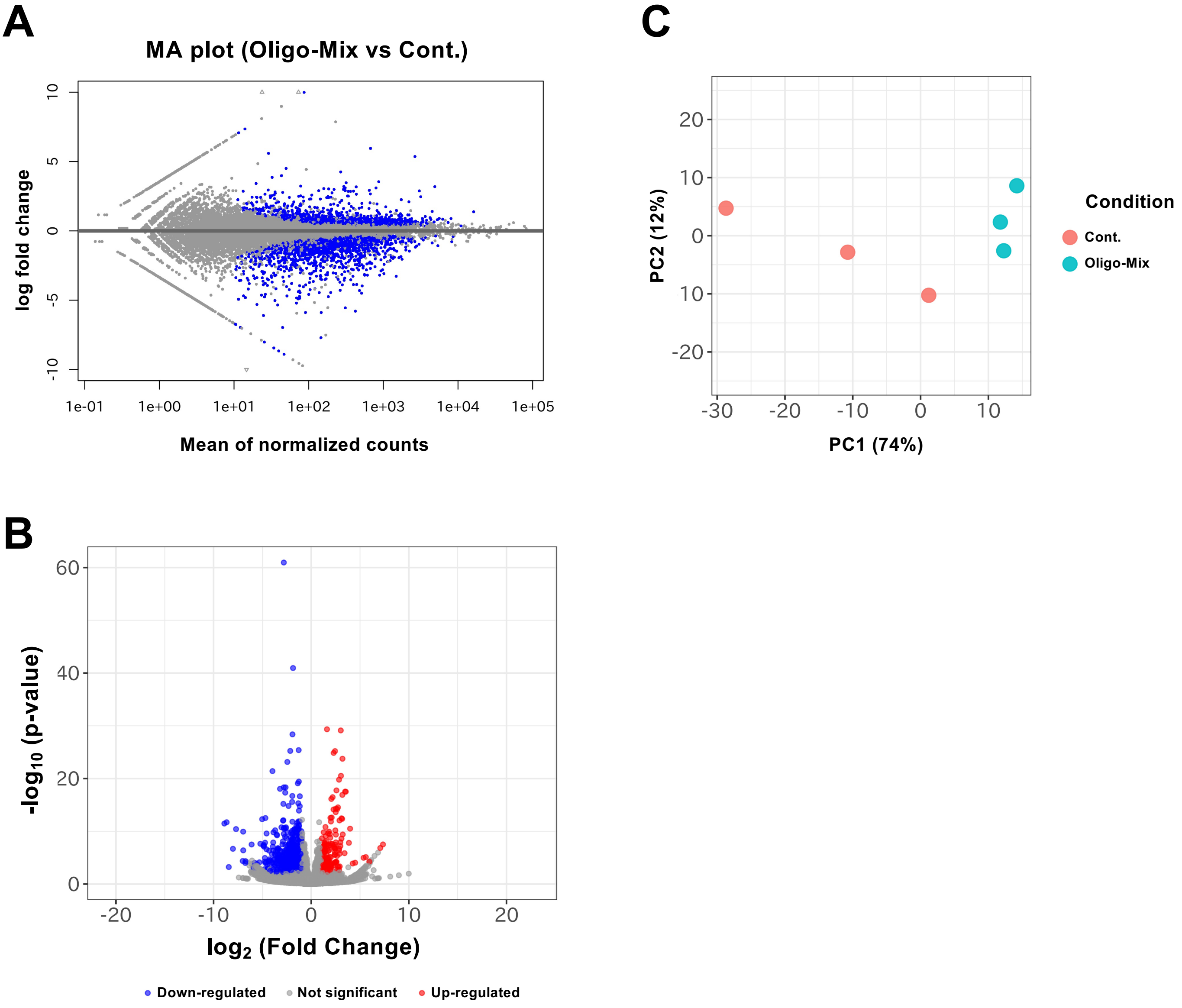


**Fig. S1** RNAseq analysis of tomato leaves treated with Oligo-Mix.
**(A)** MA plot showing $\log_2$-fold changes versus mean normalized counts for genes in Oligo-Mix treated leaves compared with controls (Cont.). Upregulated genes ($\log_2$FC > 1, FDR < 0.05) are highlighted in blue. **(B)** Volcano plot illustrating statistical significance ($-\log_{10}$ P value) against $\log_2$ fold change. Significantly upregulated and downregulated genes are shown in red and blue, respectively. **(C)** Principal component analysis (PCA) of global gene expression profiles showing separation between control and Oligo-Mix samples along PC1 (74%), indicating a transcriptional shift induced by Oligo-Mix.
