## Supplementary material for "Foliar application of mixed DAMP- and MAMP-oligosaccharides enhances root growth and fruit yield in okra and tomato in greenhouse experiments": Table S1

**Table S1**. Tomato genes upregulated by Oligo-Mix treatment (log2FC > 2, FDR < 0.05, baseMean > 10), ordered by transcript abundance (TPM) in Oligo-Mix–treated leaves.

| Gene Name / ID | Cont. (Average TPM) | Oligo-Mix (Average TPM) | baseMean | $log_2$FC (Oligo-Mix /Cont.) | *p* value | *FDR* | Protein ID | Description |
|---|---|---|---|---|---|---|---|---|
| gad2 | 1506.21 | 3586.98 | 16196 | 1.38 | 9.9E-07 | 6.0E-05 | NP_001233775.1 | 2-oxoglutarate-dependent dioxygenase 2 |
| rbcL/LyesC2p061 | 152.31 | 1337.02 | 4894 | 3.19 | 1.2E-17 | 9.0E-15 | YP_008563096.1 | Ribulose bisphosphate carboxylase large chain |
| yfe37 | 621.85 | 1507.99 | 4520 | 1.40 | 2.3E-07 | 1.7E-05 | NP_001233856.1 | Alcohol dehydrogenase |
| LOC101244831 | 281.91 | 528.13 | 3639 | 1.04 | 2.9E-03 | 3.0E-02 | NP_001306251.1 | Sterol side chain reductase 2 |
| LOC544001 | 267.60 | 1313.80 | 3171 | 2.46 | 2.2E-07 | 1.7E-05 | NP_001233769.1 | Miraculin-like precursor |
| mcpi | 62.19 | 2260.01 | 2647 | 5.36 | 1.0E-05 | 4.3E-04 | NP_001233934.1 | Metallocarboxypeptidase inhibitor precursor |
| LOC100736434 | 184.45 | 378.07 | 2213 | 1.15 | 8.3E-06 | 3.6E-04 | NP_001234494.1 | Cytochrome P450 |
| ycf1/LyesC2p009 | 33.08 | 135.46 | 2198 | 2.10 | 2.4E-13 | 1.1E-10 | YP_008563147.1 | Hypothetical chloroplast RF1 (chloroplast) |
| LOC138349000 | 164.91 | 339.50 | 2122 | 1.15 | 5.5E-06 | 2.5E-04 | XP_069154862.1 | Cytochrome P450 CYP72A616-like |
| GAME6 | 141.93 | 279.27 | 1766 | 1.10 | 1.3E-03 | 1.6E-02 | NP_001352908.1 | Glycoalkaloid metabolism 6 |
| C5-SD | 200.74 | 429.42 | 1693 | 1.23 | 2.9E-04 | 5.3E-03 | NP_001352901.1 | Delta(7)-sterol-C5(6)-desaturase |
| LOC101265706 | 123.03 | 243.40 | 1601 | 1.11 | 2.5E-03 | 2.6E-02 | XP_004251744.1 | Zeatin O-glucosyltransferase |
| SMO4 | 180.78 | 464.10 | 1501 | 1.47 | 8.6E-10 | 1.4E-07 | NP_001353020.1 | 4-methylsterol oxidase |
| LOC101248132 | 125.19 | 249.67 | 1282 | 1.10 | 2.2E-09 | 3.3E-07 | XP_004247137.1 | uncharacterized protein |
| GAME1 | 92.97 | 220.39 | 1242 | 1.37 | 2.5E-04 | 4.7E-03 | NP_001233853.2 | UDP-galactosyltransferase precursor |
| LOC101268254 | 83.45 | 170.37 | 1100 | 1.16 | 7.7E-04 | 1.1E-02 | XP_004251001.1 | Zeatin O-glucosyltransferase |
| GAME18 | 54.98 | 167.05 | 1076 | 1.74 | 1.1E-05 | 4.3E-04 | XP_004243636.2 | beta-D-glucosyl crocetin beta-1,6-glucosyltransferase |
| SMO3 | 117.19 | 213.60 | 958 | 1.00 | 5.2E-04 | 8.4E-03 | NP_001353008.1 | Sterol methyl oxidase 3 |
| LOC100147717 | 164.28 | 360.48 | 893 | 1.22 | 2.0E-04 | 4.0E-03 | XP_004240704.2 | Protein PELPK2 |
| psbT/LyesC2p043 | 298.19 | 3325.65 | 868 | 3.54 | 3.1E-18 | 2.4E-15 | YP_008563115.1 | Photosystem II protein T (chloroplast) |
| HMGS1 | 54.80 | 124.95 | 858 | 1.32 | 2.5E-04 | 4.7E-03 | XP_004245661.1 | Hydroxymethylglutaryl-CoA synthase |
| LOC101254427 | 80.10 | 204.03 | 825 | 1.46 | 1.5E-11 | 4.3E-09 | XP_004248391.2 | Inositol oxygenase 1 |
| CEVI-1 | 64.46 | 228.54 | 823 | 1.94 | 2.3E-04 | 4.4E-03 | NP_001234132.2 | Peroxidase precursor |
| HYD1 | 124.35 | 203.10 | 817 | 1.10 | 4.8E-05 | 1.4E-03 | NP_001353019.1 | 3-beta-hydroxysteroid-Delta(8),Delta(7)-isomerase |
| LOC101257156 | 21.91 | 172.76 | 797 | 3.10 | 2.4E-09 | 3.6E-07 | XP_069148497.1 | UDP-glycosyltransferase 75C1-like |
| LOC101267346 | 30.05 | 58.57 | 684 | 1.08 | 2.1E-05 | 7.5E-04 | XP_010322950.1 | Phosphate dikinase, chloroplastic |
| TD2 | 2.33 | 135.26 | 673 | 5.95 | 5.9E-05 | 1.7E-03 | NP_001296095.1 | Threonine dehydratase 2 biosynthetic, chloroplastic |
| CHS2 | 29.29 | 171.17 | 670 | 2.67 | 6.2E-15 | 3.0E-12 | NP_001234036.2 | Chalcone synthase 2 |
| atpF/LyesC2p083 | 106.87 | 412.56 | 663 | 2.02 | 1.4E-12 | 4.7E-10 | YP_008563074.1 | ATP synthase $CF_0$ subunit I (chloroplast) |
| AnthOMT | 36.09 | 250.40 | 650 | 2.92 | 4.6E-04 | 7.5E-03 | NP_001289828.1 | Flavonoid 3',5'-methyltransferase |
| matK/LyesC2p088 | 46.95 | 134.80 | 643 | 1.61 | 4.6E-30 | 2.6E-26 | YP_008563069.1 | Maturase K (chloroplast) |
| LOC101268074 | 110.70 | 215.53 | 623 | 1.06 | 3.8E-05 | 1.2E-03 | XP_004252262.1 | H/ACA ribonucleoprotein complex subunit 2-like protein |
| LOC101264429 | 72.66 | 140.46 | 556 | 1.04 | 8.1E-05 | 2.1E-03 | XP_004240482.1 | Multiple organellar RNA editing factor 2, chloroplastic |
| LOC101261260 | 44.05 | 135.78 | 517 | 1.70 | 4.5E-05 | 1.3E-03 | XP_004228671.1 | Lignin-forming anionic peroxidase |
| LOC101250474 | 20.97 | 79.43 | 517 | 1.98 | 2.2E-08 | 2.6E-06 | XP_010316283.1 | Benzaldehyde dehydrogenase, mitochondrial |
| LOC101244055 | 112.28 | 229.48 | 500 | 1.15 | 9.5E-06 | 4.0E-04 | NP_001309349.1 | CDGSH iron-sulfur domain-containing protein |
| LOC101259187 | 39.73 | 111.64 | 475 | 1.63 | 8.0E-05 | 2.1E-03 | XP_004246615.2 | Xyloglucan endotransglucosylase/hydrolase protein 32 |
| LOC101261977 | 34.81 | 77.13 | 452 | 1.27 | 2.7E-06 | 1.4E-04 | XP_004249621.1 | UDP-glycosyltransferase 86A1-like |
| LOC101264884 | 69.52 | 141.58 | 448 | 1.15 | 3.6E-07 | 2.5E-05 | XP_069147503.1 | Heavy metal-associated isoprenylated plant protein 7-like |
| ANS | 17.31 | 97.83 | 435 | 2.65 | 1.9E-03 | 2.1E-02 | XP_019070824.1 | Anthocyanidin synthase |
| rps14/LyesC2p071 | 63.84 | 559.56 | 433 | 3.17 | 4.4E-13 | 1.7E-10 | YP_008563086.1 | Ribosomal protein S14 (chloroplast) |
| LOC101246821 | 53.59 | 120.25 | 432 | 1.28 | 1.6E-10 | 3.2E-08 | XP_004237884.1 | FCS-Like Zinc finger 5 |
| LOC101264938 | 30.30 | 59.85 | 430 | 1.07 | 8.4E-05 | 2.2E-03 | XP_004241450.1 | DEK domain-containing chromatin-associated protein 1 |
| LOC100191129 | 13.59 | 90.22 | 399 | 2.86 | 1.9E-07 | 1.5E-05 | NP_001234419.1 | Anthocyanin acyltransferase |
| DWF5-2 | 26.49 | 76.03 | 398 | 1.63 | 2.5E-07 | 1.9E-05 | NP_001353043.1 | 7-dehydrocholesterol reductase |
| LOC101250395 | 24.29 | 75.62 | 371 | 1.76 | 5.1E-07 | 3.4E-05 | XP_004237444.2 | Stress enhanced protein 2, chloroplastic |
| LOC138338147 | 2.79 | 19.11 | 363 | 2.85 | 1.6E-20 | 2.0E-17 | XP_069144959.1 | Cytochrome b6-f complex subunit 4 |
| LOC101262648 | 26.46 | 67.00 | 357 | 1.45 | 2.2E-05 | 7.6E-04 | XP_004246628.1 | Heavy metal-associated isoprenylated plant protein 3 |
| CHI1 | 21.72 | 114.49 | 340 | 2.51 | 2.4E-14 | 1.0E-11 | NP_001307640.1 | Chalcone--flavonone isomerase |
| DFR | 11.51 | 75.37 | 340 | 2.85 | 4.2E-07 | 2.9E-05 | NP_001234408.2 | Dihydroflavonol 4-reductase |
| JRE4 | 23.68 | 125.37 | 331 | 2.52 | 5.1E-15 | 2.5E-12 | XP_004229751.1 | Ethylene-responsive transcription factor ERF098 |
| MKB1 | 67.92 | 132.33 | 330 | 1.09 | 3.6E-03 | 3.4E-02 | NP_001352941.1 | RING-finger E3 ubiquitin ligase Makibishi 1 |
| GS1 | 28.32 | 66.27 | 324 | 1.34 | 3.8E-08 | 3.9E-06 | NP_001306784.1 | Glutamine synthetase |
| LOC101248118 | 8.91 | 94.93 | 318 | 3.47 | 2.9E-18 | 2.4E-15 | XP_004242780.2 | uncharacterized protein |
| LOC101265881 | 13.32 | 90.42 | 310 | 2.90 | 1.9E-06 | 1.0E-04 | XP_004232669.1 | Glutathione S-transferase F11 |
| LOC101244316 | 7.73 | 62.05 | 308 | 3.10 | 3.5E-13 | 1.4E-10 | XP_004247056.1 | Anthocyanidin-3-O-glucoside rhamnosyltransferase |
| rps19/LyesC2p029 | 47.10 | 412.82 | 296 | 3.20 | 1.8E-24 | 3.1E-21 | YP_008563129.1 | Ribosomal protein S19 (chloroplast) |
| LOC543899 | 8.95 | 64.67 | 293 | 2.99 | 6.2E-08 | 5.9E-06 | NP_001234424.1 | uncharacterized protein/ Multi antimicrobial extrusion family |
| LOC101266223 | 33.13 | 95.44 | 292 | 1.61 | 1.0E-05 | 4.2E-04 | XP_010321455.1 | Chalcone isomerase-like protein 2 |
| LOC101260642 | 80.00 | 159.91 | 286 | 1.10 | 1.3E-03 | 1.6E-02 | XP_004239779.1 | Histone H4 |
| LOC101262948 | 32.91 | 83.09 | 277 | 1.35 | 8.2E-07 | 5.1E-05 | - | uncharacterized lncRNA |
| rpl2/LyesC2p028 | 39.30 | 143.31 | 277 | 2.19 | 3.4E-17 | 2.2E-14 | YP_008563130.1 | Ribosomal protein L2 (chloroplast) |
| LOC101259314 | 29.69 | 62.48 | 271 | 1.17 | 1.2E-04 | 2.8E-03 | XP_004234117.1 | uncharacterized protein/ PDDEXK-like family |
| LOC101247557 | 12.30 | 216.30 | 269 | 4.25 | 1.3E-04 | 2.9E-03 | NP_001315434.1 | Proteinase inhibitor I precursor |
| LOC101262552 | 35.09 | 69.00 | 261 | 1.08 | 1.6E-03 | 1.9E-02 | XP_004246712.2 | Glutathione S-transferase L3 |
| SFP5 | 18.80 | 41.96 | 254 | 1.29 | 2.4E-05 | 8.2E-04 | XP_025887547.1 | Sugar-porter family protein 5 |
| LOC101258850 | 16.40 | 32.09 | 252 | 1.09 | 1.0E-04 | 2.5E-03 | XP_004237077.1 | NRT1/ PTR family 1.2 |
| LOC101248301 | 7.72 | 25.04 | 246 | 1.84 | 3.6E-07 | 2.5E-05 | XP_004240219.1 | Zeatin O-xylosyltransferase-like |
| LOC101247008 | 35.11 | 73.28 | 240 | 1.17 | 1.2E-03 | 1.6E-02 | XP_004236870.2 | High mobility group B protein 7 |
| LOC104649465 | 50.39 | 112.29 | 237 | 1.25 | 4.9E-03 | 4.4E-02 | XP_010327073.1 | uncharacterized protein |
| LOC101251423 | 23.06 | 55.81 | 235 | 1.38 | 2.1E-08 | 2.5E-06 | XP_004248667.1 | uncharacterized protein/MIP aquaporin family |
| F3'5'H | 6.42 | 45.87 | 218 | 2.99 | 1.2E-08 | 1.5E-06 | NP_001234840.2 | Flavonoid 3'5' hydroxylase |
| LOC101262509 | 17.73 | 63.55 | 216 | 1.93 | 7.2E-05 | 1.9E-03 | XP_004237402.1 | PLAT domain-containing protein 2 |
| LOC101259456 | 11.20 | 20.72 | 210 | 1.01 | 1.4E-03 | 1.7E-02 | XP_004243487.1 | Cellulose synthase-like protein H1 |
| F3H | 9.59 | 59.96 | 205 | 2.74 | 2.9E-15 | 1.5E-12 | NP_001316412.1 | Flavanone 3-dioxygenase |
| LOC101260668 | 21.37 | 59.32 | 203 | 1.57 | 2.9E-06 | 1.5E-04 | XP_004246621.1 | uncharacterized protein |
| LOC101248277 | 17.13 | 45.29 | 201 | 1.48 | 4.3E-04 | 7.1E-03 | XP_004235206.1 | Superoxide dismutase [Fe], chloroplastic |
| GA20OX1 | 18.47 | 35.68 | 199 | 1.03 | 1.3E-03 | 1.6E-02 | NP_001234070.1 | Gibberellin 20-oxidase-1 |
| rpoA/LyesC2p038 | 13.78 | 70.59 | 198 | 2.44 | 6.2E-26 | 1.4E-22 | YP_008563120.1 | RNA polymerase alpha subunit (chloroplast) |
| LOC101266025 | 45.23 | 116.30 | 197 | 1.42 | 8.6E-04 | 1.2E-02 | NP_001317094.1 | Gibberellin-regulated family protein precursor |
| LOC101266234 | 22.54 | 77.41 | 197 | 1.90 | 7.4E-06 | 3.2E-04 | - | uncharacterized lncRNA |
| LOC101255991 | 17.43 | 36.67 | 196 | 1.18 | 1.8E-05 | 6.6E-04 | XP_010320821.1 | UDP-glycosyltransferase 89A2 |
| LOC101251684 | 28.35 | 62.65 | 196 | 1.23 | 8.2E-08 | 7.5E-06 | XP_004240522.1 | Glutaredoxin-C13 |
| ODC | 14.77 | 37.53 | 192 | 1.44 | 5.1E-08 | 5.0E-06 | NP_001234616.1 | Ornithine decarboxylase |
| LOC101255179 | 17.54 | 43.12 | 191 | 1.35 | 8.8E-08 | 8.0E-06 | XP_004234958.2 | Zinc finger protein CONSTANS-LIKE 10 |
| LOC101255495 | 29.81 | 66.91 | 185 | 1.26 | 9.3E-04 | 1.3E-02 | XP_004240096.2 | Heavy metal-associated isoprenylated plant protein 39-like |
| rpl22/LyesC2p030 | 19.62 | 150.80 | 185 | 3.03 | 7.5E-30 | 3.3E-26 | YP_008563128.1 | Ribosomal protein L22 (chloroplast) |
| CHS1 | 9.52 | 45.98 | 184 | 2.39 | 2.0E-07 | 1.6E-05 | NP_001234033.2 | Chalcone synthase 1 |
| trnS-GCU/LyesC2t005 | 140.64 | 754.78 | 182 | 2.47 | 6.7E-11 | 1.5E-08 | - | tRNA-Ser (chloroplast) |
| LOC101265837 | 11.75 | 31.18 | 180 | 1.51 | 8.1E-09 | 1.1E-06 | XP_069154960.1 | Reticulon-like protein B2 |
| LOC101244151 | 23.92 | 47.06 | 179 | 1.06 | 7.5E-06 | 3.3E-04 | XP_010316929.1 | Urease accessory protein G-like |
| LOC104646221 | 8.72 | 45.37 | 174 | 2.40 | 2.5E-06 | 1.3E-04 | - | uncharacterized lncRNA |
| LOC101261090 | 19.28 | 37.80 | 173 | 1.06 | 2.7E-03 | 2.8E-02 | XP_004233250.1 | Pyrroline-5-carboxylate reductase |
| ycf3/LyesC2p068 | 15.28 | 120.87 | 159 | 3.05 | 3.0E-21 | 4.0E-18 | YP_008563089.1 | Photosystem I assembly protein ycf3 (chloroplast) |
| LOC100750250 | 17.27 | 50.66 | 157 | 1.63 | 4.0E-05 | 1.2E-03 | NP_001238797.2 | Short-chain dehydrogenase-reductase |

| | | | | | | | | |
|---|---|---|---|---|---|---|---|---|
| LOC101244167 | 63.14 | 154.96 | 157 | 1.37 | 2.0E-05 | 7.0E-04 | XP_004235193.2 | Glycine-rich cell wall structural protein 2 |
| LOC101265488 | 6.05 | 14.88 | 153 | 1.39 | 2.8E-07 | 2.1E-05 | XP_004248713.1 | Isoflavone reductase homolog |
| LOC101265469 | 14.20 | 28.72 | 152 | 1.10 | 1.6E-03 | 1.9E-02 | XP_004246551.1 | Ankyrin repeat domain-containing protein EMB506, chloroplastic |
| LOC101261737 | 12.35 | 30.11 | 146 | 1.37 | 1.3E-08 | 1.6E-06 | NP_001311389.1 | Alpha/beta-Hydrolases superfamily protein precursor |
| rps3/LyesC2p031 | 16.76 | 76.64 | 143 | 2.29 | 1.4E-25 | 2.7E-22 | YP_008563127.1 | Ribosomal protein S3 (chloroplast) |
| LOC101257827 | 9.19 | 21.00 | 139 | 1.26 | 9.5E-04 | 1.3E-02 | XP_004232727.2 | RNA pseudouridine synthase 6, chloroplastic |
| LOC104647298 | 28.61 | 57.36 | 138 | 1.09 | 6.0E-05 | 1.7E-03 | XP_069153909.1 | uncharacterized protein |
| LOC101260093 | 4.84 | 34.08 | 136 | 2.90 | 8.6E-13 | 3.1E-10 | XP_004249447.1 | Anthocyanidin 3-O-glucosyltransferase |
| LOC101255624 | 12.02 | 24.59 | 134 | 1.14 | 1.8E-04 | 3.7E-03 | XP_004247084.1 | uncharacterized protein/SNARE associated protein |
| ZIP | 12.84 | 32.98 | 132 | 1.46 | 8.4E-04 | 1.2E-02 | NP_001233865.1 | bZIP transcription factor |
| RDRP | 5.21 | 10.26 | 132 | 1.04 | 1.5E-03 | 1.8E-02 | NP_001234319.1 | RNA-directed RNA polymerase |
| LOC101247117 | 7.70 | 22.81 | 130 | 1.67 | 4.8E-04 | 7.8E-03 | XP_004253114.2 | Protein PIN-LIKES 3-like |
| LOC109120423 | 14.55 | 28.87 | 128 | 1.10 | 3.8E-04 | 6.6E-03 | XP_019069696.1 | uncharacterized protein |
| LOC101258706 | 7.94 | 16.44 | 126 | 1.12 | 3.1E-04 | 5.5E-03 | XP_004248692.1 | DETOXIFICATION 46, chloroplastic-like |
| LOC101258933 | 6.71 | 20.28 | 121 | 1.70 | 8.9E-04 | 1.3E-02 | XP_004236064.1 | Cytochrome P450 78A5-like |
| LOC101259237 | 10.16 | 25.07 | 116 | 1.37 | 1.0E-03 | 1.3E-02 | XP_004237470.1 | Protein SLOW GREEN 1, chloroplastic |
| LOC101255053 | 17.74 | 34.34 | 115 | 1.08 | 7.4E-05 | 2.0E-03 | XP_069147763.1 | Acyl carrier protein 3, mitochondrial |
| WHY1 | 13.04 | 28.09 | 115 | 1.18 | 1.2E-03 | 1.6E-02 | NP_001289829.2 | DNA-binding protein WHY1 |
| LOC101253959 | 5.58 | 32.01 | 110 | 2.58 | 1.8E-18 | 1.5E-15 | XP_004232713.1 | Senescence regulator protein S40-5 |
| XTH6 | 8.42 | 38.34 | 107 | 2.30 | 1.0E-07 | 9.2E-06 | NP_001233819.2 | Xyloglucan endotransglucosylase-hydrolase XTH6 |
| LOC101255228 | 13.40 | 31.69 | 106 | 1.35 | 6.5E-05 | 1.8E-03 | XP_004245848.2 | FCS-Like Zinc finger 1 |
| LOC101248084 | 8.92 | 16.39 | 105 | 1.01 | 1.8E-03 | 2.1E-02 | - | probable pseudogene (7-deoxyloganetin glucosyltransferase-like) |
| LOC101247128 | 11.46 | 35.60 | 103 | 1.73 | 2.4E-10 | 4.6E-08 | XP_004240422.1 | uncharacterized protein |
| rps4/LyesC2p067 | 12.87 | 60.01 | 103 | 2.29 | 7.2E-15 | 3.4E-12 | YP_008563090.1 | Ribosomal protein S4 (chloroplast) |
| LOC101262376 | 10.59 | 21.39 | 101 | 1.10 | 1.7E-05 | 6.2E-04 | XP_004233007.1 | GDSL esterase/lipase At1g29670-like |
| LOC104649457 | 4.98 | 10.23 | 100 | 1.16 | 1.9E-03 | 2.1E-02 | XP_010327017.1 | uncharacterized protein |
| LOC101256307 | 7.60 | 15.83 | 99 | 1.17 | 3.7E-05 | 1.2E-03 | XP_004243643.1 | Scarecrow-like protein 23 |
| LOC101252727 | 12.85 | 31.21 | 95 | 1.37 | 3.5E-05 | 1.1E-03 | XP_004230652.1 | Peptide methionine sulfoxide reductase B5 |
| LOC101266842 | 6.18 | 14.46 | 95 | 1.32 | 5.1E-04 | 8.2E-03 | XP_004244012.1 | Amino acid permease 7 |
| LOC101249848 | 4.02 | 10.44 | 91 | 1.46 | 3.2E-03 | 3.2E-02 | XP_004246412.1 | Cation/H(+) antiporter 20 |
| LOC101253549 | 12.21 | 26.43 | 90 | 1.17 | 3.0E-03 | 3.0E-02 | XP_069147457.1 | GRF1-interacting factor 1 |
| LOC101260003 | 8.63 | 17.51 | 87 | 1.12 | 2.1E-04 | 4.1E-03 | XP_004232649.1 | Zinc transporter 3 |
| LOC101253231 | 7.89 | 16.24 | 87 | 1.14 | 3.4E-03 | 3.3E-02 | XP_004230743.1 | Polygalacturonase inhibitor 1 |
| LOC101259699 | 8.09 | 18.03 | 86 | 1.24 | 6.4E-05 | 1.8E-03 | XP_004232736.1 | CRIB domain-containing protein RIC10 |
| atp1 | 4.67 | 19.72 | 86 | 2.15 | 2.4E-08 | 2.7E-06 | YP_009430460.1 | ATP synthase subunit 1 (mitochondrion) |
| LOC101257261 | 2.16 | 11.28 | 84 | 2.44 | 3.4E-08 | 3.7E-06 | XP_004238938.1 | NRT1/ PTR family protein 3.1 |
| LOC101259820 | 12.00 | 25.05 | 84 | 1.15 | 9.4E-05 | 2.3E-03 | XP_004235161.1 | Peroxisomal membrane protein 11B |
| LOC101257731 | 17.59 | 37.20 | 82 | 1.17 | 7.5E-04 | 1.1E-02 | XP_010317149.1 | STS14 protein |
| GID1b-2 | 6.47 | 13.58 | 79 | 1.16 | 4.0E-03 | 3.8E-02 | NP_001352577.1 | Gibberellin receptor GID1b-2 |
| LOC101258712 | 12.09 | 25.62 | 79 | 1.18 | 1.5E-03 | 1.8E-02 | XP_004249524.1 | Organelle RRM domain-containing protein 2, mitochondrial |
| LOC101247471 | 13.45 | 43.58 | 79 | 1.84 | 3.0E-03 | 3.1E-02 | XP_004230980.1 | Protein RADIALIS-like 4 |
| LOC101265216 | 3.93 | 7.71 | 78 | 1.06 | 2.7E-03 | 2.8E-02 | XP_004235676.1 | Homeobox-leucine zipper protein GLABRA 2 |
| LOC101245519 | 3.29 | 11.95 | 78 | 1.95 | 1.7E-09 | 2.6E-07 | XP_004232860.1 | Allantoinase |
| LOC101253552 | 3.95 | 36.61 | 78 | 3.23 | 4.0E-10 | 7.1E-08 | XP_004250256.1 | Mavicyanin-like |
| LOC101253378 | 14.88 | 30.42 | 77 | 1.13 | 1.0E-04 | 2.5E-03 | XP_069153450.1 | uncharacterized protein At5g64816-like |
| LOC101258746 | 4.93 | 12.54 | 76 | 1.42 | 6.4E-07 | 4.1E-05 | XP_004237675.1 | uncharacterized protein/Amino acid permease |
| LOC101259773 | 5.38 | 11.76 | 76 | 1.23 | 6.5E-05 | 1.8E-03 | XP_004245100.1 | Zinc transporter 4, chloroplastic |
| LOC101261415 | 4.00 | 10.78 | 75 | 1.54 | 2.3E-04 | 4.5E-03 | XP_004240117.2 | Pectinesterase/pectinesterase inhibitor 12 |
| LOC138341709 | 2.00 | 4.53 | 74 | 1.26 | 4.3E-04 | 7.2E-03 | - | uncharacterized lncRNA |
| cuAO | 2.00 | 10.91 | 74 | 2.50 | 6.3E-05 | 1.8E-03 | NP_001296994.1 | Copper amine oxidase |
| LOC101247356 | 6.29 | 17.97 | 72 | 1.61 | 2.0E-03 | 2.2E-02 | XP_004246214.1 | Acylsugar acyltransferase 3-like |
| LOC101256520 | 6.88 | 15.45 | 72 | 1.25 | 5.8E-04 | 9.1E-03 | XP_004247602.1 | Perakine reductase |
| LOC101266674 | 6.18 | 13.80 | 71 | 1.25 | 2.0E-03 | 2.2E-02 | XP_004248176.2 | Aminotransferase TAT2 |
| TCP23 | 6.85 | 13.33 | 70 | 1.09 | 5.2E-03 | 4.5E-02 | NP_001386546.1 | TCP transcription factor 23 |
| LOC101248091 | 4.52 | 9.27 | 70 | 1.11 | 4.6E-03 | 4.2E-02 | XP_004236624.1 | Remorin 4.2-like |
| LOC101259420 | 4.72 | 12.01 | 70 | 1.44 | 2.6E-04 | 4.8E-03 | XP_004252985.1 | Adenosylhomocysteinase |
| LOC101248444 | 5.65 | 12.61 | 70 | 1.24 | 1.3E-04 | 3.0E-03 | XP_004249986.1 | Serine/threonine-protein kinase D6PKL2-like |
| LOC101266953 | 5.58 | 12.66 | 68 | 1.28 | 4.3E-05 | 1.3E-03 | XP_004242673.2 | Stemmadenine O-acetyltransferase |
| LOC138338601 | 0.98 | 2.94 | 68 | 1.68 | 3.2E-08 | 3.4E-06 | - | probable pseudogene (protein Ycf2-like) |
| LOC101253442 | 6.00 | 13.25 | 67 | 1.20 | 4.1E-03 | 3.8E-02 | XP_004232389.1 | uncharacterized protein |
| LOC112941069 | 3.71 | 7.65 | 64 | 1.13 | 2.5E-04 | 4.7E-03 | XP_025885664.1 | IRK-interacting protein-like |
| LOC104646389 | 16.34 | 31.38 | 64 | 1.04 | 1.3E-03 | 1.6E-02 | XP_010318049.2 | Senescence regulator protein S40-7 |
| LOC101261573 | 6.26 | 11.94 | 64 | 1.00 | 2.6E-04 | 4.9E-03 | XP_004233085.1 | Purine permease 10 |
| LE16 | 8.46 | 36.33 | 63 | 2.09 | 5.3E-04 | 8.5E-03 | NP_001233953.1 | Non-specific lipid-transfer protein 2 |
| LOC101244564 | 5.04 | 10.57 | 62 | 1.13 | 9.6E-04 | 1.3E-02 | NP_001334360.1 | Serine carboxypeptidase family protein |
| trnM-CAU/LyesC2t026 | 111.42 | 249.97 | 61 | 1.23 | 2.5E-04 | 4.7E-03 | - | tRNA-Met (chloroplast) |
| LOC101247253 | 10.07 | 25.11 | 60 | 1.40 | 1.2E-04 | 2.8E-03 | XP_004229337.2 | Non-specific lipid transfer protein GPI-anchored 14 |
| LOC101262396 | 5.76 | 15.62 | 60 | 1.50 | 1.4E-03 | 1.7E-02 | XP_010314505.2 | Calcineurin B-like protein 4 |
| LOC101267105 | 2.93 | 9.65 | 59 | 1.79 | 2.8E-05 | 9.4E-04 | XP_004236685.1 | Ethylene-response factor C3-like |
| LOC101262086 | 8.03 | 16.34 | 56 | 1.11 | 3.3E-04 | 5.9E-03 | XP_004252243.1 | Single-stranded DNA-binding protein, mitochondrial |
| LOC101250736 | 3.08 | 5.88 | 55 | 1.01 | 1.7E-03 | 2.0E-02 | XP_004229807.1 | LRR receptor kinase BAK1 |
| LOC138339710 | 8.89 | 22.27 | 53 | 1.41 | 2.0E-04 | 4.0E-03 | XP_069147752.1 | Non-specific lipid transfer protein GPI-anchored 14-like |
| LOC101250593 | 11.18 | 23.74 | 53 | 1.17 | 1.6E-03 | 1.9E-02 | XP_004236796.1 | Leucine zipper transcription factors, c-Myc-binding |
| LOC112941068 | 5.00 | 15.30 | 51 | 1.66 | 4.2E-03 | 3.9E-02 | - | uncharacterized lncRNA |
| LOC101248997 | 4.20 | 8.83 | 50 | 1.13 | 3.2E-03 | 3.2E-02 | XP_004245491.1 | Protein ABIL2-like |
| cathDInh | 1.05 | 25.61 | 50 | 4.50 | 9.0E-05 | 2.3E-03 | NP_001234142.2 | Cathepsin D inhibitor protein |
| trnR-UCU/LyesC2t008 | 87.00 | 201.55 | 49 | 1.30 | 6.1E-04 | 9.4E-03 | - | tRNA-Arg (chloroplast) |
| LOC101258621 | 1.44 | 2.80 | 48 | 1.04 | 3.8E-03 | 3.6E-02 | XP_004233864.1 | Pentatricopeptide repeat-containing protein, mitochondrial |
| LOC101264324 | 5.31 | 11.98 | 48 | 1.24 | 1.9E-03 | 2.1E-02 | XP_004238792.1 | Origin of replication complex subunit 6 |
| LOC101256047 | 3.60 | 9.03 | 48 | 1.40 | 1.6E-05 | 6.0E-04 | XP_069146337.1 | Cycloartenol-C-24-methyltransferase-like |
| LOC101264159 | 4.05 | 9.94 | 48 | 1.40 | 4.2E-04 | 7.0E-03 | XP_004248082.1 | E3 ubiquitin-protein ligase SINAT3 |
| LOC101246275 | 5.74 | 20.46 | 47 | 1.93 | 2.3E-03 | 2.4E-02 | XP_004246743.1 | uncharacterized protein |
| rpl23/LyesC2p027 | 15.65 | 57.14 | 47 | 1.83 | 4.5E-08 | 4.6E-06 | YP_008563131.1 | Ribosomal protein L23 (chloroplast) |
| LOC101249461 | 1.42 | 3.63 | 47 | 1.47 | 1.7E-03 | 1.9E-02 | - | probable pseudogene (Metal-nicotianamine transporter YSL7) |
| LOC101249176 | 3.32 | 8.68 | 47 | 1.51 | 2.3E-03 | 2.4E-02 | XP_069154516.1 | Zeatin O-xylosyltransferase-like |
| LOC101260211 | 3.87 | 9.11 | 47 | 1.32 | 1.4E-03 | 1.7E-02 | XP_025885695.2 | 7-deoxyloganetic acid glucosyltransferase |
| LOC101268864 | 2.40 | 6.23 | 46 | 1.46 | 1.8E-05 | 6.6E-04 | XP_004238976.1 | Piriformospora indica-insensitive protein 2 |
| LOC101256061 | 5.56 | 11.22 | 45 | 1.09 | 3.0E-03 | 3.0E-02 | XP_004251289.1 | Chitinase 2-like |
| LOC138339597 | 9.39 | 22.71 | 45 | 1.35 | 5.8E-04 | 9.1E-03 | - | uncharacterized lncRNA |
| LOC101261729 | 4.61 | 8.80 | 44 | 1.01 | 2.5E-03 | 2.7E-02 | XP_010324378.1 | D-amino-acid transaminase, chloroplastic-like |
| LOC101256734 | 4.04 | 8.24 | 43 | 1.11 | 1.3E-03 | 1.6E-02 | XP_004250011.1 | uncharacterized protein |
| LOC101259115 | 1.01 | 15.80 | 43 | 3.98 | 3.1E-11 | 8.1E-09 | XP_025884886.2 | RNA exonuclease 4-like |
| LOC104648558 | 1.59 | 4.44 | 42 | 1.58 | 3.2E-04 | 5.8E-03 | XP_069143660.1 | G-type lectin S-receptor-like serine/threonine-protein kinase |
| LOC101259352 | 2.39 | 7.62 | 41 | 1.70 | 1.3E-03 | 1.6E-02 | NP_001315978.1 | Methyltransferase PrmA-like |
| LOC101252681 | 1.94 | 5.50 | 41 | 1.58 | 2.8E-04 | 5.1E-03 | XP_025886627.1 | Growth-regulating factor 1-like |
| LOC101254527 | 1.25 | 5.06 | 40 | 2.09 | 1.8E-04 | 3.7E-03 | XP_010326368.2 | Transcription factor BHLH42 |
| LOC101258206 | 7.45 | 20.44 | 39 | 1.51 | 1.8E-05 | 6.7E-04 | XP_004248984.1 | Large ribosomal subunit protein eL43 |
| LOC101244203 | 5.04 | 12.70 | 38 | 1.42 | 5.7E-04 | 9.0E-03 | XP_004243685.1 | uncharacterized protein |
| LecRK-IV | 2.38 | 4.61 | 38 | 1.04 | 5.0E-03 | 4.5E-02 | NP_001307189.1 | L-type lectin-domain containing receptor kinase IV.1-like |

| | | | | | | | | |
|---|---|---|---|---|---|---|---|---|
| LOC101258797 | 1.48 | 5.85 | 37 | 2.01 | 7.9E-06 | 3.4E-04 | - | uncharacterized lncRNA |
| rps18/LyesC2p047 | 13.65 | 37.30 | 36 | 1.51 | 1.6E-04 | 3.5E-03 | YP_008563110.1 | Ribosomal protein S18 (chloroplast) |
| psbI/LyesC2p085 | 45.08 | 92.83 | 36 | 1.15 | 4.8E-03 | 4.3E-02 | YP_008563072.1 | Photosystem II protein I (chloroplast) |
| LOC101252820 | 1.12 | 7.10 | 36 | 2.72 | 4.4E-04 | 7.3E-03 | XP_004247901.2 | 5-OH-xanthotoxin synthase |
| LOC101264620 | 9.61 | 63.61 | 36 | 2.81 | 6.7E-04 | 1.0E-02 | - | uncharacterized lncRNA |
| LOC101268385 | 3.34 | 9.03 | 35 | 1.53 | 6.0E-04 | 9.4E-03 | - | uncharacterized lncRNA |
| LOC101255514 | 2.83 | 13.82 | 35 | 2.32 | 2.6E-04 | 4.9E-03 | XP_004244317.2 | uncharacterized protein/SOUL heme-binding protein |
| LOC101244049 | 1.50 | 4.43 | 35 | 1.62 | 1.8E-04 | 3.8E-03 | XP_004231946.1 | Growth-regulating factor 1-like |
| LOC101245813 | 1.15 | 2.86 | 35 | 1.42 | 5.9E-04 | 9.2E-03 | XP_069146162.1 | Wall-associated receptor kinase 17-like |
| LOG8 | 3.57 | 21.32 | 35 | 2.59 | 6.2E-05 | 1.7E-03 | NP_001244912.1 | Cytokinin riboside 5'-monophosphate phosphoribohydrolase LOG8 |
| LOC101267773 | 2.38 | 7.86 | 34 | 1.80 | 4.8E-03 | 4.3E-02 | XP_004233955.1 | uncharacterized protein |
| LOC138338255 | 0.76 | 1.58 | 33 | 1.13 | 5.5E-03 | 4.7E-02 | - | probable pseudogene |
| LOC101265642 | 3.61 | 7.76 | 33 | 1.18 | 1.8E-03 | 2.0E-02 | XP_004239629.1 | Protein OPI10 homolog |
| MYB75 | 2.37 | 13.24 | 32 | 2.55 | 6.4E-05 | 1.8E-03 | NP_001265992.1 | Transcription factor MYB75 |
| LOC101251649 | 0.72 | 10.87 | 32 | 3.86 | 1.5E-08 | 1.9E-06 | XP_004233702.2 | RNA exonuclease 4 |
| LOC101260516 | 2.44 | 5.99 | 32 | 1.34 | 4.3E-03 | 4.0E-02 | XP_004235056.1 | uncharacterized protein |
| LOC109120460 | 2.27 | 6.77 | 31 | 1.65 | 1.1E-04 | 2.7E-03 | - | uncharacterized lncRNA |
| LOC104644842 | 1.07 | 6.29 | 31 | 2.41 | 9.0E-05 | 2.3E-03 | - | uncharacterized lncRNA |
| LOC101248064 | 3.24 | 10.08 | 30 | 1.66 | 3.4E-05 | 1.1E-03 | - | uncharacterized lncRNA |
| LOC101251470 | 0.08 | 4.18 | 29 | 5.59 | 7.7E-06 | 3.3E-04 | XP_004238404.1 | uncharacterized protein/S9A serine peptidase family |
| LOC101255707 | 3.57 | 8.98 | 29 | 1.39 | 8.2E-04 | 1.2E-02 | XP_004242010.1 | uncharacterized protein/Serine hydrolase (FSH1) |
| trnT-UGU/LyesC2t018 | 39.83 | 126.76 | 28 | 1.72 | 4.5E-05 | 1.3E-03 | - | tRNA-Thr (chloroplast) |
| psbZ/LyesC2p072 | 13.62 | 47.74 | 27 | 1.83 | 3.6E-04 | 6.3E-03 | YP_008563085.1 | Photosystem II protein Z (chloroplast) |
| LOC101267002 | 1.28 | 4.01 | 27 | 1.69 | 5.3E-03 | 4.6E-02 | XP_004235076.2 | Amino acid transporter AVT1A |
| LOC101245157 | 2.08 | 7.30 | 27 | 1.87 | 9.2E-05 | 2.3E-03 | NP_001333113.1 | Isoaspartyl peptidase/L-asparaginase 2 |
| trnH-GUG/LyesC2t001 | 33.29 | 122.83 | 27 | 1.95 | 5.6E-04 | 8.9E-03 | - | tRNA-His (chloroplast) |
| LOC101253408 | 0.57 | 3.83 | 26 | 2.65 | 4.6E-06 | 2.1E-04 | XP_004245759.1 | uncharacterized protein |
| LOC101264349 | 2.56 | 6.80 | 25 | 1.45 | 2.5E-03 | 2.7E-02 | XP_004244728.1 | MYB-like transcription factor 4 |
| LOC101259901 | 3.60 | 10.82 | 25 | 1.67 | 1.6E-03 | 1.9E-02 | XP_004232327.1 | Cyclin-dependent protein kinase inhibitor SMR6 |
| LOC101260499 | 2.05 | 6.01 | 24 | 1.57 | 5.7E-04 | 9.0E-03 | XP_004250193.1 | Nudix hydrolase 18, mitochondrial-like |
| LOC101247045 | 1.97 | 5.41 | 24 | 1.56 | 5.2E-03 | 4.6E-02 | XP_019070612.1 | Cell number regulator 1-like |
| trnR-ACG/LyesC2t036 | 20.03 | 108.13 | 23 | 2.42 | 9.2E-07 | 5.7E-05 | - | tRNA-Arg (chloroplast) |
| LOC101256615 | 0.93 | 2.53 | 22 | 1.46 | 1.4E-03 | 1.8E-02 | XP_004228489.1 | Scarecrow-like protein 3 |
| ycf2/LyesC2p004 | 0.33 | 0.94 | 21 | 1.55 | 1.3E-03 | 1.6E-02 | YP_008563132.1 | Hypothetical chloroplast RF2 (chloroplast) |
| LOC101262483 | 2.23 | 6.39 | 21 | 1.55 | 5.2E-03 | 4.6E-02 | XP_004250781.1 | LOB domain-containing protein 4 |
| MYB | 1.29 | 7.42 | 20 | 2.41 | 9.3E-05 | 2.3E-03 | NP_001234262.1 | R2R3MYB transcription factor 14 |
| LOC138348360 | 0.67 | 2.83 | 20 | 2.08 | 5.3E-03 | 4.6E-02 | XP_004238437.1 | Serine/threonine-protein kinase PBL21 |
| LOC101263989 | 1.03 | 3.01 | 20 | 1.61 | 3.2E-03 | 3.2E-02 | XP_004253086.2 | Growth-regulating factor 4 |
| psbN/LyesC2p042 | 5.52 | 58.59 | 20 | 3.41 | 1.4E-06 | 7.9E-05 | YP_008563116.1 | Photosystem II protein N |
| accD/LyesC2p060 | 1.30 | 3.71 | 18 | 1.52 | 3.3E-03 | 3.2E-02 | YP_008563097.1 | Acetyl-CoA carboxylase carboxyltransferase β subunit (chloroplast) |
| LOC101257801 | 1.20 | 5.09 | 17 | 2.01 | 1.6E-03 | 1.9E-02 | XP_004246880.2 | NAC domain-containing protein 90-like |
| rpl16/LyesC2p032 | 3.26 | 12.30 | 15 | 1.94 | 1.9E-03 | 2.1E-02 | YP_008563126.1 | Ribosomal protein L16 (chloroplast) |
| LOC101267009 | 0.77 | 3.18 | 15 | 1.98 | 4.0E-03 | 3.8E-02 | XP_004253011.1 | uncharacterized protein |
| LOC101263191 | 0.00 | 5.74 | 14 | 7.35 | 3.1E-08 | 3.4E-06 | XP_004252057.2 | Protein ABIL5 |
| rpl32/LyesC2p019 | 8.13 | 26.18 | 14 | 1.67 | 5.0E-03 | 4.4E-02 | YP_008563137.1 | Ribosomal protein L32 (chloroplast) |
| LOC101268240 | 0.37 | 2.98 | 13 | 2.86 | 2.3E-03 | 2.4E-02 | XP_004249643.3 | UDP-glycosyltransferase 76E1-like |
| LOC109119841 | 0.00 | 3.98 | 12 | 7.07 | 1.4E-07 | 1.3E-05 | XP_019068534.1 | Bidirectional sugar transporter N3-like |
